# Proper chromosome alignment depends on BRCA2 phosphorylation by PLK1

**DOI:** 10.1101/265934

**Authors:** Åsa Ehlén, Charlotte Martin, Simona Miron, Manon Julien, François-Xavier Theillet, Virginie Ropars, Gaetana Sessa, Romane Beaurepere, Virginie Boucherit, Patricia Duchambon, Ahmed El Marjou, Sophie Zinn-Justin, Aura Carreira

## Abstract

The BRCA2 tumor suppressor protein is involved in the maintenance of genome integrity through its role in homologous recombination. In mitosis, BRCA2 is phosphorylated by Polo-like kinase 1 (PLK1). Here we describe how this phosphorylation contributes to the control of mitosis. We identified two highly conserved phosphorylation sites at S193 and T207 of BRCA2. Phosphorylated-T207 is a *bona fide* docking site for PLK1 as illustrated by the crystal structure of the BRCA2 peptide bound to PLK1 Polo-box domain. We found that BRCA2 bound to PLK1 forms a complex with the phosphatase PP2A and phosphorylated-BUBR1. Reducing BRCA2 binding to PLK1, as observed in *BRCA2* breast cancer variants S206C and T207A, alters the tetrameric complex resulting in misaligned chromosomes, faulty chromosome segregation and aneuploidy. We thus reveal a direct role of BRCA2 in the alignment of chromosomes, distinct from its DNA repair function, with important consequences on chromosome stability. These findings may explain in part the aneuploidy observed in *BRCA2*-mutated tumors.

## Introduction

The BRCA2 tumor suppressor protein plays an important role in DNA repair by homologous recombination (HR) ^1, 2^ that takes place preferentially during S/G2 phases of the cell cycle ^3^. BRCA2 has also emerging functions in mitosis, for example, at the kinetochore, it forms a complex with BUBR1 ^4, 5^, an essential factor for the faithful segregation of chromosomes that is required for kinetochore-microtubule attachment and is a component of the spindle assembly checkpoint (SAC) ^6, 7^. These two activities of BUBR1 involve different partners and are functionally distinct ^8, 9^. BRCA2 has been proposed to contribute to BUBR1 SAC activity through direct binding and promoting its acetylation by the acetyl transferase PCAF ^4, 10^, although due to confounding results in the BUBR1 interaction site in BRCA2, it is unclear if this interaction is direct ^4, 5^. At the end of mitosis, BRCA2 localizes to the midbody and assists cell division by serving as a scaffold protein for the central spindle components ^11–13^. In mitosis, BRCA2 is a target of phosphorylation by PLK1 both in its N-terminal region ^14 15^ and in its central region ^15^, although the functional role of these phosphorylation events remains unclear.

PLK1 is a master regulator of the cell cycle that is upregulated in mitosis ^16, 17^. Among other functions, PLK1 directly binds and phosphorylates BUBR1 at several residues including the attachment-sensitive site S670 ^18^ and the two tension-sensitive sites S676 ^19^ and T680 ^20^ in prometaphase allowing the formation of stable kinetochore-microtubule attachments. This activity needs to be tightly regulated to ensure proper alignment of the chromosomes at the metaphase plate ^8, 9, 19^. The kinase activity of Aurora B is an essential to destabilize erroneous kinetochore-microtubule interactions ^21^ whereas the phosphatase PP2A protects initial kinetochore-microtubule interactions from excessive destabilization by Aurora B ^22^. This function is achieved through the interaction of PP2A-B56 subunit with BUBR1 phosphorylated at the Kinetochore Attachment and Regulatory Domain (KARD) motif (including residues S670, S676 and T680) ^20^. Thus, the interplay between PLK1, BUBR1, Aurora B and PP2A is necessary for the formation of stable kinetochore-microtubule attachments.

PLK1 is recruited to specific targets via its Polo-box domain (PBD) ^23^. PBD interacts with phosphosites characterized by the consensus motif S-[pS/pT]-P/X ^24^. These phosphosites are provided by a priming phosphorylation event, usually mediated by CDK1 or other proline-directed kinases ^17^; however, there is also evidence that PLK1 itself might create docking sites (“self-priming”) during cytokinesis ^25, 26^.

Several BRCA2 sites have been suggested as phosphorylated by PLK1 in mitosis, some of which belong to a cluster of serines and threonines located in BRCA2 N-terminus around residue S193 ^14^. We set out to investigate which of these sites are phosphorylated by PLK1, and to reveal whether these phosphorylation events play a role in the regulation of mitotic progression. Here, we demonstrate that S193 and T207 are the only two residues that can be phosphorylated by PLK1 in the conserved region around S193, and we further reveal that phosphorylated BRCA2-T207 is a *bona fide* docking site for PLK1. By investigating the phenotype of BRCA2 missense variants that limit the phosphorylation of BRCA2-T207, we reveal an unexpected role for BRCA2 in the alignment of chromosomes at the metaphase plate. We demonstrate that phosphorylation of BRCA2-T207 by PLK1 facilitates the formation of a complex between BRCA2-PLK1-pBUBR1 and the phosphatase PP2A required to counter excessive Aurora B activity at the kinetochores ^20^. A defect in this function of BRCA2 manifested in chromosome misalignment, chromosome segregation errors, mitotic delay and aneuploidy, leading to chromosome instability.

## Results

### *BRCA2* variants identified in breast cancer reduce the PLK1-dependent phosphorylation of BRCA2 N-terminal region

Several missense variants of uncertain significance (VUS) identified in *BRCA2* in breast cancer patients are located in the N-terminal region predicted to be phosphorylated by PLK1 (around S193) (Breast information core (BIC) ^27^ and BRCAShare^28^), summarized in Supplementary table 1. To find out if any of these variants affected PLK1 phosphorylation in this region, we purified fragments comprising amino acids 1 to 250 of BRCA2 (hereafter BRCA2_1-250_) from human embryonic kidney cells (HEK293T) and used an *in vitro* kinase assay to assess the phosphorylation by PLK1 of the fragments containing either the WT sequence, the different BRCA2 variants M192T, S196N, S206C and T207A, or the mutant S193A, previously reported to reduce the phosphorylation of BRCA2 by PLK1 ^14^. As expected, S193A reduced the phosphorylation of BRCA2_1-250_ by PLK1 (Fig. 1a, 1b). Interestingly, variants T207A and S206C also led to a 2-fold decrease in PLK1 phosphorylation of BRCA2_1-250_ (Fig. 1a, b). In contrast, M192T slightly increased the phosphorylation above WT levels whereas S196N did not significantly modify the phosphorylation of BRCA2_1-250_ by PLK1 (Fig. 1a, b). The phosphorylation observed in the BRCA2 fragments is specific of the recombinant PLK1 kinase as it is PLK1 concentration dependent (Supplementary Fig. 1a, b) and when replacing the PLK1-WT by a kinase-dead (PLK1-KD) version of the protein (K82R) ^29^, purified using the same protocol, or adding a PLK1 inhibitor (BI2536) to the reaction, the phosphorylation of BRCA2_1-250_ decreases significantly (Fig. 1c, lanes 4 and 5 compared to lane 3; Fig. 1d).

**Figure 1.**
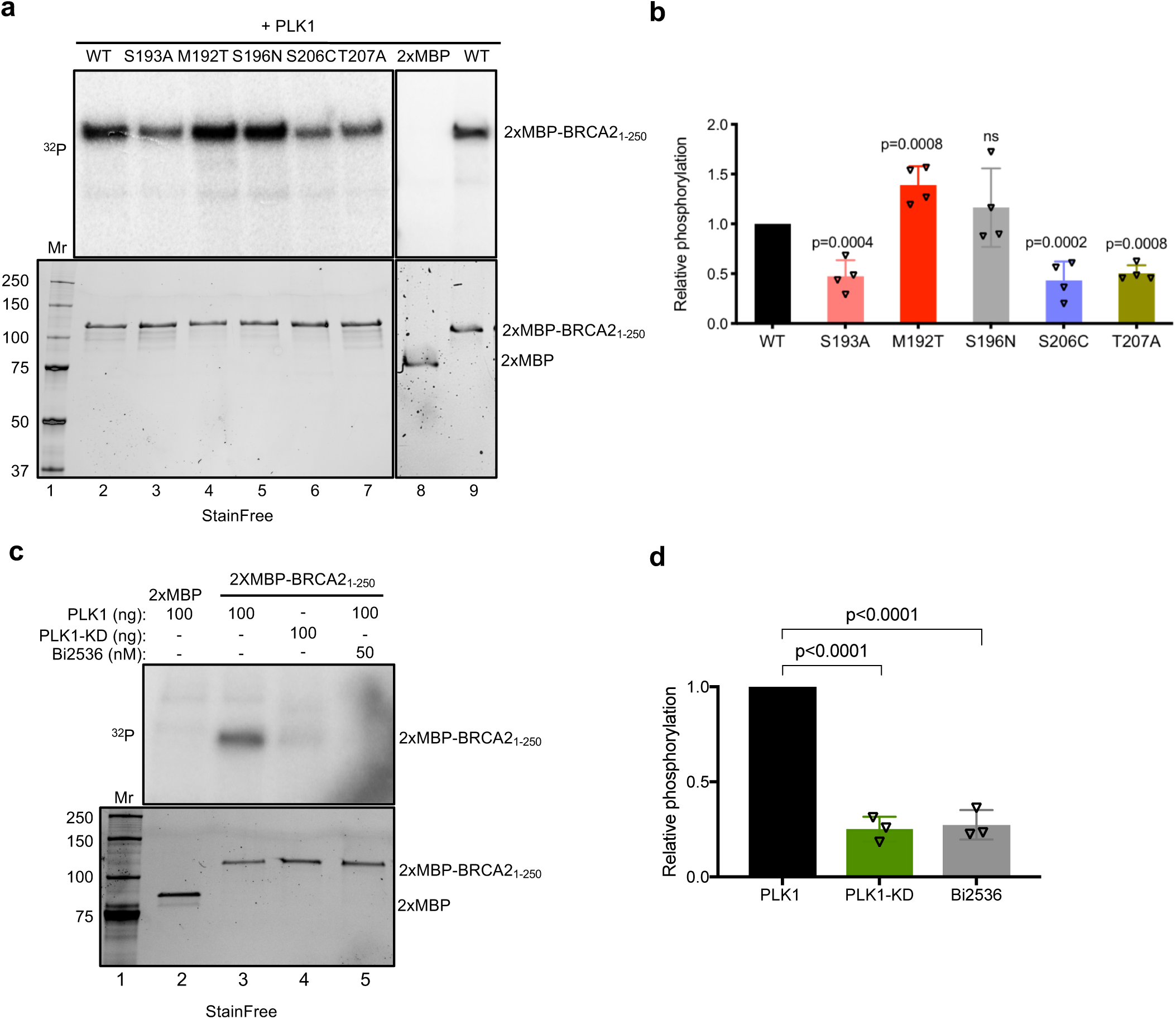
BRCA2 VUS alter PLK1 phosphorylation of BRCA2_1-250_. **(a)** PLK1 *in vitro* kinase assay with BRCA2_1-250_. Top: The polypeptides encompassing 2x-MBP-BRCA2_1-250_ WT or S193A, M192T, S196N, T200K, S206C, T207A mutations or the 2XMBP tag were incubated with recombinant PLK1 in the presence of γ^32^P-ATP. The samples were resolved on 7.5% SDS-PAGE and the ^32^P-labeled products were detected by autoradiography. Bottom: 7.5% SDS-PAGE showing the input of purified 2xMBP-BRCA2_1-_ 250 WT and mutated proteins (0.5 μg) used in the reaction as indicated. **(b)** Quantification of the relative phosphorylation in (A). Data in (B) are represented as mean ± SD from at least four independent experiments (WT (n=9), S193A (n=4), M192T (n=4), S196N (n=4), S206C (n=4), T200K (n=5), T207A (n=4)). **(c)** PLK1 *in vitro* kinase assay performed as in (A) with recombinant PLK1 or the PLK1 kinase dead K82R mutant (PLK1-KD) together with BRCA2_1-250_ WT as substrate, in the presence or absence of the PLK1 inhibitor BI2536 (50 nM) in the kinase reaction buffer. **(d)** Quantification of the relative phosphorylation in (c). Data in (d) are represented as mean ± SD from three independent experiments. (b and c) One-way ANOVA test with Dunnett’s multiple comparisons test was used to calculate statistical significance of differences (the p-values show differences compared to WT (b) or PLK1 (d); ns (non-significant)).

Together, these results show that VUS T207A and S206C identified in breast cancer patients impair phosphorylation of BRCA2_1-250_ by PLK1 *in vitro*.

### BRCA2-T207 is a target of phosphorylation by PLK1

The reduction of BRCA2 phosphorylation in BRCA2_1-250_ containing T207A and S206C variants suggested that these residues could be targets for PLK1 phosphorylation. We investigated this possibility by following the PLK1 phosphorylation kinetics of two overlapping fragments of BRCA2 N-terminus comprising S206 and T207 (hereafter BRCA2_48-218_ and BRCA2_190-284_) using Nuclear Magnetic Resonance (NMR) spectroscopy (Fig. 2a). Together, these fragments cover a large N-terminal region of BRCA2 including the cluster of conserved residues around S193 (from amino acid 180 to amino acid 210; Supplementary Fig.1c). NMR analysis allows residue-specific quantification of ^15^N-labelled peptide phosphorylation. Fig. 2b shows superimposed ^1^H-^15^N HSQC spectra of BRCA2_48-218_ and BRCA2_190-284_ before (black) and after (red) phosphorylation with recombinant PLK1. Analysis of these experiments revealed phosphorylation of S193 and of three other phosphosites, including T207, by PLK1 in the BRCA2 region from amino acid 48 to amino acid 284. Interestingly, while T219 and T226 conservation is poor, T207 and S193 are conserved from mammals to fishes (Supplementary Fig. 1c) suggesting that both residues are important for BRCA2 function.

**Figure 2.**
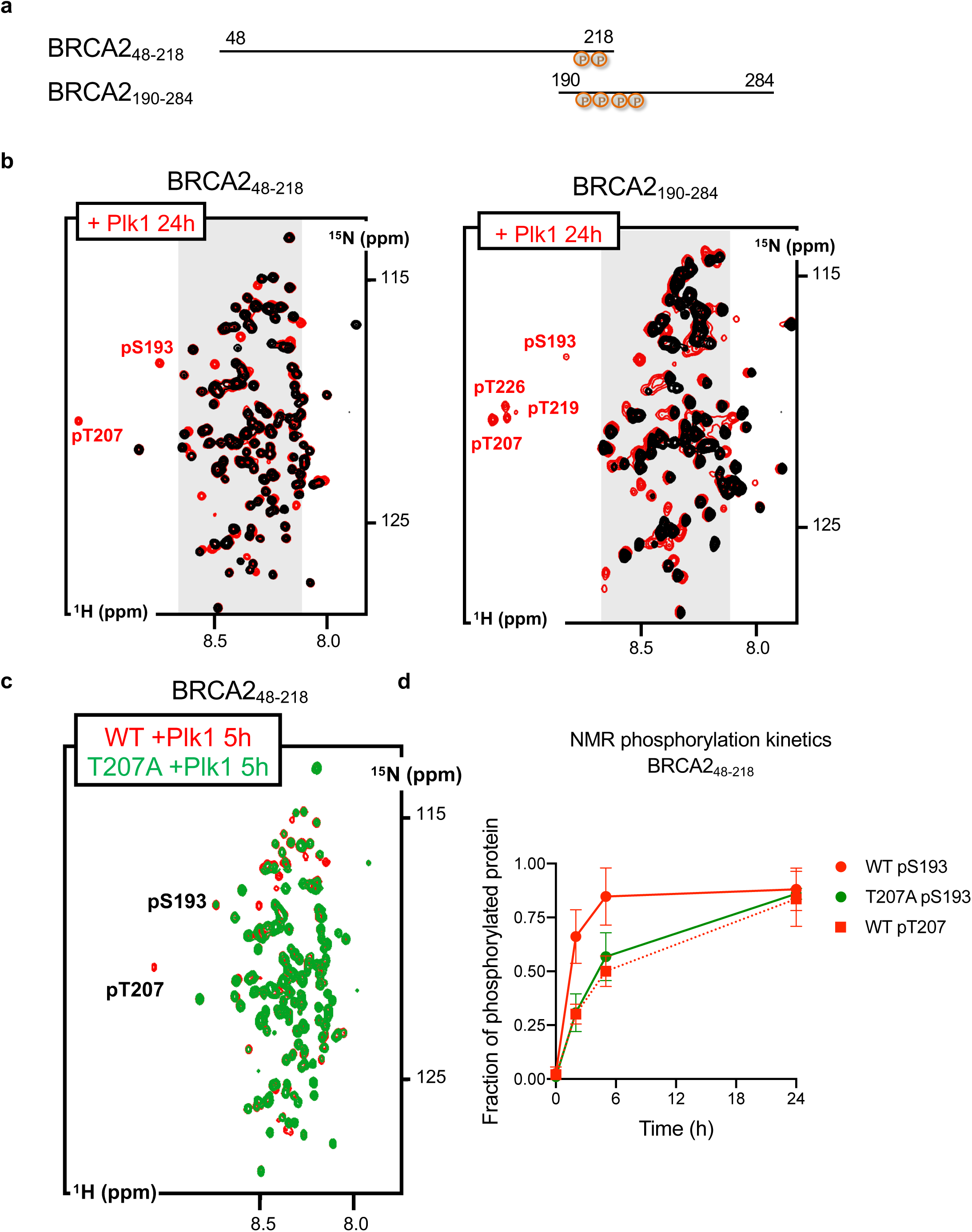
PLK1 phosphorylates T207 in BRCA2_48-218_ and BRCA2_190-284_. Phosphorylation of BRCA2_48-218_ and BRCA2_190-284_ by PLK1 as observed by NMR. **(a)** Schematic view of the 2 overlapping BRCA2 fragments analysed by NMR. Residues identified as phosphorylated in (b) are indicated. **(b)** Superposition of the ^1^H-^15^N HSQC spectra recorded before (black) and after (24h; red) incubation with PLK1. Each spectrum contains one peak per backbone NH group. Most peaks are located in the grey region of the spectrum, indicating that they correspond to disordered residues. Assignment of each peak to a BRCA2 residue was performed using a classical heteronuclear NMR strategy. Peaks corresponding to phosphorylated residues are indicated. **(c)** Comparison of the phosphorylation kinetics recorded for BRCA2_48-218_ WT and T207A. ^1^H-^15^N HSQC spectra of BRCA2_48-218_WT (red) and T207A (green) recorded 5 h after addition of PLK1 are superimposed to highlight the overall decrease of phosphorylation observed for the mutant compared to the WT. **(d)** Fraction of phosphorylated protein deduced from the intensities of the peaks corresponding to the non-phosphorylated and phosphorylated residues plotted as a function of time. WT S193 and T207 time points are represented by red circles and squares, respectively, while T207A S193 timepoints are represented by green circles. The graph represents results of three independent experiments; error bars (SD).

### BRCA2 variant T207A alters the phosphorylation kinetics by PLK1

Having identified T207 as a target of phosphorylation of PLK1, we next compared the residue-specific phosphorylation kinetics in the polypeptide WT BRCA2_48-218_ containing the variant T207A that displayed reduced overall phosphorylation (Fig. 1a, 1b). (The production of a ^15^N-labelled recombinant fragment comprising S206C yielded an insoluble protein precluding NMR analysis). Time-resolved NMR experiments revealed that PLK1 phosphorylates significantly less BRCA2_48-218_ containing the variant T207A than the WT peptide (Fig. 2c). The initial phosphorylation rate of S193 was decreased by a factor of 2 (Fig. 2d), and T207 was as expected not phosphorylated, being mutated into an alanine. Similar results were obtained using BRCA2_190-284_ (Supplementary Fig. 2). This NMR analysis is in agreement with the *in vitro* kinase assay performed using the BRCA2_1-250_ fragment purified from human cells (Fig. 1), in which T207A reduces the phosphorylation of BRCA2_1-250_ fragment by PLK1.

### Variants T207A and S206C reduce the interaction of BRCA2 and PLK1

The finding that T207 is efficiently phosphorylated by PLK1 in BRCA2_48-218_ and BRCA2_190-_284 (Fig. 2b) together with the observation that T207A mutation causes a global decrease in the phosphorylation of these fragments (Fig. 2c; Supplementary Fig. 2) and the prediction that T207 is a docking site for PLK1_PBD_ binding ^24^ made us hypothesize that T207 might be a “self-priming” phosphorylation event required for the interaction of PLK1 with BRCA2 at this site. If so, the variants that reduce phosphorylation of T207 by PLK1 would be predicted to alter PLK1_PBD_ binding. To test this hypothesis, we examined the interaction of PLK1 with the VUS-containing polypeptides. We overexpressed 2xMBP-BRCA2_1-250_ constructs carrying these variants in U2OS cells to detect the endogenous PLK1 that co-immunoprecipitates with 2xMBP-BRCA2_1-250_ using amylose pull-down. As expected, overexpressed BRCA2_1-250_ was able to interact with endogenous PLK1 from mitotic cells but not from asynchronous cells (predominantly in G1/S) where the levels of PLK1 are reduced (Fig. 3a, lane 2 compared to lane 1). Furthermore, the variants T207A and S206C showed a weaker interaction with PLK1 than the WT protein (Fig. 3a, pull-down lanes 4, 6 compared to lane 2, Fig. 3b) despite the protein levels of PLK1 remaining unchanged (Fig. 3a, compare PLK1 input lanes 4 and 6 to lane 2). In contrast, the effect of M192T and S196N on the interaction was mild (Fig. 3a, compare pull-down lanes 10 and 12 to lane 8, Fig. 3b). These results are consistent with the idea of a self-priming phosphorylation by PLK1 on T207.

**Figure 3.**
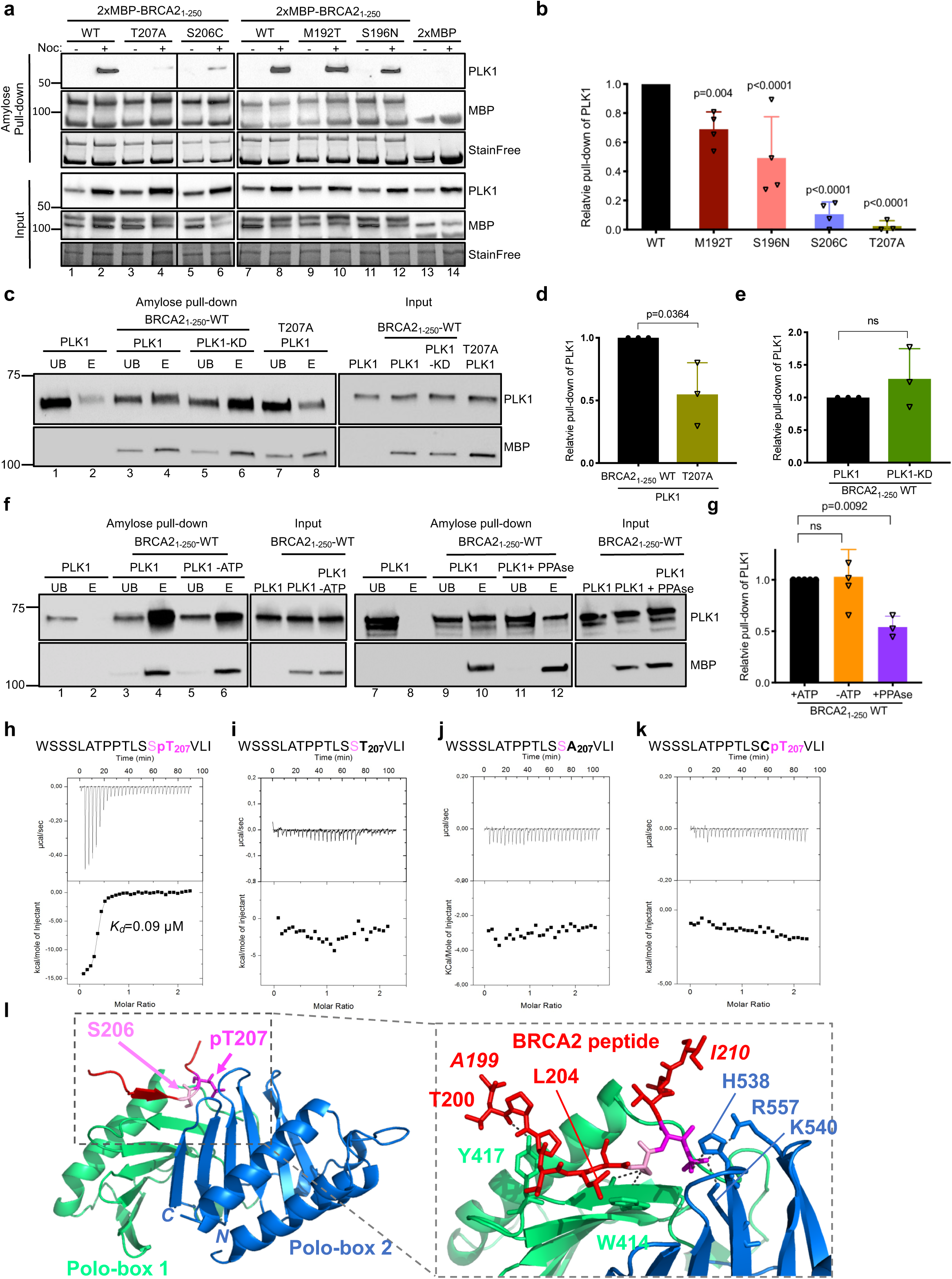
BRCA2 variants showing reduced phosphorylation by PLK1 impair PLK1 binding. **(a)** 2xMBP-BRCA2_1-250_ expressing the WT, the variants (M192T, S196N, S206C and T207A) and the 2XMBP-tag were expressed in U2OS cells by transient transfection for 30h before the cells were treated with nocodazole for 14h. Mitotic cells were lysed and immunoprecipitation was performed against the MBP tag using amylose beads. Complexes were resolved on 4-15% SDS-PAGE followed by western blotting using anti-PLK1 and anti-MBP antibodies. StainFree images of the gels before transfer was used as loading control (cropped image is shown). **(b)** Quantification of co-immunoprecipitated PLK1 with 2xMBP-BRCA2_1-250_ in (a), relative to the input levels of PLK1. Results are presented as the fold change compared to BRCA2_1-250_-WT. The data represents the mean ± SD of 3-4 independent experiments (WT (n=4), M192T (n=4), S196N (n=4), S206C (n=4), T207A (n=3)). Statistical significance of the difference in between WT and the variants was calculated with one-way ANOVA test with Dunnett’s multiple comparisons test. **(c)** PLK1 *in vitro* kinase assay followed by *in vitro* pull-down assay using amylose beads. The polypeptides 2x-MBP-BRCA2_1-250_ WT and T207A were incubated with recombinant PLK1-WT or the kinase dead K82R mutant (PLK1-KD, only for the BRCA2_1-250_ –WT fragment) in kinase buffer with ATP followed by incubation with amylose beads. The bound complexes were eluted from the beads with maltose and resolved on 10% SDS-PAGE followed by western blotting using anti-PLK1 and anti-MBP antibodies. UB: unbound fraction, E: maltose-eluted fraction. **(d-e)** Quantification of the PLK1 pull-down in (c) relative to the PLK1 levels in the input. Results are presented as the fold change compared to BRCA2_1-250_-WT in (d) and PLK1-WT in (e). The data represents the mean ± SD of three independent experiments. **(f)** PLK1 *in vitro* kinase assay followed by *in vitro* pull-down assay using amylose beads as in (c). *Left panel:* The 2x-MBP-BRCA2_1-250_-WT polypeptide was incubated with recombinant PLK1-WT in the absence or presence of ATP followed by incubation with amylose beads. *Right panel:* The 2x-MBP-BRCA2_1-250_-WT polypeptide was pre-treated with phosphatase (FastAP Thermosensitive Alkaline Phosphatase) for 1 hour before the incubation with recombinant PLK1-WT followed by the amylose-beads incubation as described in (c). **(g)** Quantification of the PLK1 pull-down in (f) relative to the PLK1 levels in the input. Results are presented as the fold change compared to kinase assay performed with non-phosphatase treated BRCA2_1-250_-WT in the presence of ATP. The data represents the mean ± SD of three-four independent experiments. Statistical significance of the difference in (d-e, g) was calculated with two-tailed t-test (the p-values show significant differences; ns (non-significant)) **(h-k)** Isothermal Titration Calorimetry (ITC) thermograms showing binding of PLK1_PBD_ to a 17 aa BRCA2 peptide containing **(h)** pT207, **(i)** T207, **(j)** A207, **(k)** C206pT207. Residues S206 and pT207 are highlighted in pink (S206) and magenta (pT207) in the peptide sequences. **(l)** 3D cartoon representation of the crystal structure of the interface between PLK1_PBD_ (Polo-box 1 in green and Polo-box 2 in blue) and the BRCA2 peptide containing pT207 (in red except for S206 and pT207 that are highlighted in pink and magenta, respectively, as in (h-i)). The left panel shows a global view of the structure, whereas the right panel focuses on the interface between PLK1_PBD_ and the BRCA2 peptide (from A199 to I210, marked in italics). In the left panel, S206 and pT207 are represented in sticks, whereas in the right panel, all the BRCA2 residues observed in the crystal structure are displayed in sticks. The hydrogen bonds involving side chain atoms are displayed and depicted as dark grey dots. The amino acids of PLK1_PBD_ involved in these interactions are highlighted in sticks representation.

To provide further evidence that the PLK1-mediated phosphorylation of BRCA2 favors BRCA2 binding, we performed an *in vitro* kinase assay with recombinant proteins followed by an amylose pull-down and eluted the bound proteins with maltose. PLK1 was found in the maltose elution with BRCA2_1-250_-WT demonstrating that PLK1-phosphorylated BRCA2_1-250_ binds to PLK1 (Fig. 3c lane 4, Fig. 3d). In contrast, the fraction of PLK1 in the eluate of BRCA2_1-250_-T207A was substantially reduced (Fig. 3c, lane 8 compared to lane 4, Fig. 3d) indicating that the phosphorylation of T207 is required for efficient binding to PLK1 and confirming our results with cell lysates (Fig. 3a, b). Strikingly, we observed no difference in PLK1 binding between BRCA2_1-250_-WT phosphorylated by PLK1 or its kinase dead mutant (PLK1-KD; Fig. 3c, lane 6 compared to lane 4, Fig. 3e) or in the absence of ATP (Fig. 3f, lane 6 compared to lane 4, Fig. 3g). However, pre-incubating BRCA2_1-250_-WT with phosphatase before the addition of PLK1 resulted in a 2-fold decrease in the binding to PLK1 indicating that the phosphorylation of BRCA2 is required for the interaction with PLK1 (Fig. 3f lane 12 compared to 10, Fig. 3g).

### T207 is a *bona fide* docking site for PLK1

To directly demonstrate the recognition of pT207 by PLK1, we measured the affinity of recombinant PLK1_PBD_ (the target recognition domain of PLK1) for a synthetic 17 aa peptide comprising phosphorylated T207. Using isothermal titration calorimetry (ITC), we found that recombinant PLK1_PBD_ bound to the T207 phosphorylated peptide with an affinity of K*_d_*= 0.09 ± 0.01 µM (Fig. 3h), similar to the optimal affinity reported for an interaction between PLK1_PBD_ and its phosphorylated target ^24^. Consistently, PLK1_PBD_ bound to the fragment BRCA2_190-284_ with nanomolar affinity upon phosphorylation by PLK1 (K*_d_*= 0.14 ± 0.02 µM; Supplementary Fig. 3a), whereas it did not bind to the corresponding non-phosphorylated polypeptides (Fig. 3i, Supplementary Fig. 3b). Mutation T207A also abolished the interaction (Fig. 3j), in agreement with the pull-down experiments (Fig. 3a-d). More surprisingly, the phosphomimetic substitution T207D was not sufficient to create a binding site for PLK_PDB_ (Supplementary Fig. 2c). A peptide comprising pT207 and the mutation S206C could not bind to PLK1_PBD_ (Fig. 3k), as predicted from the consensus sequence requirement for PLK1_PBD_ interaction ^24^. Last, a peptide containing phosphorylated S197, which is also a predicted docking site for PLK1, bound with much less affinity to PLK1_PBD_ than pT207 (K*_d_*= 17 ± 2 µM; Supplementary Fig. 2d).

To further characterize this molecular interaction, we determined the crystal structure of PLK1_PBD_ bound to the T207 phosphorylated peptide at 3.1Å resolution (Supplementary table 2). Analysis of this 3D structure showed that, as expected, the 17 aa BRCA2 phosphopeptide binds in the cleft formed between the two Polo boxes (Fig. 3l). Twelve residues of the peptide (from A199 to I210) are well-structured upon binding, burying about 694 Å^2^ in the interface with PLK1_PBD_. BRCA2 residues from P202 to L209 are at least 25% buried in the complex. The interface between BRCA2 and PLK1_PBD_ is stabilized by 12 hydrogen bonds: the backbone of residues T200 to L209 as well as the side chain of S206 are bonded to residues from Polo Box 1, whereas the side chain of phosphorylated T207 is bonded to residues from Polo Box 2 (see the zoom view in Fig. 3l). Specifically, the backbone oxygen of T200 is bonded to the side chain of Y417 of PLK1_PBD_. Residues L204, S205 and S206 form a β-sheet with residues W414, V415 and D416, characterized by hydrogen bonds between the backbone atoms of BRCA2 L204, S206 and PLK1 D416, W414, respectively. The side chain of S206 participates in 2 hydrogen-bonding interactions with the backbone of W414, which explains the strict requirement for this amino acid at this position ^24^. Moreover, the phosphate group of pT207 participates in 3 hydrogen-bonding interactions with the side chains of residues H538, K540 and R557 in Polo Box 2 (see the zoom view in Fig. 3l). This explains the critical dependence on phosphorylation for binding observed by ITC (Fig. 3h, i). The presence of a buried leucine at the pT-3 position ^30^, as well as the electrostatic interactions of the serine at the pT-1 position with PLK1 W414 ^24^ and the phosphorylated threonine with PLK1 H538 and K540, have been described as essential for the high affinity and specificity of the interaction in other PLK1_PBD_-phosphopeptide complexes ^24, 31^.

Thus, our biochemical and structural analysis demonstrate that the BRCA2-T207 phosphopeptide interacts with PLK1_PBD_ as an optimal and specific PLK1_PBD_ ligand. It supports a mechanism in which phosphorylation of T207 by PLK1 promotes the interaction of PLK1 with BRCA2 through a *bona fide* docking site for PLK1 and favours a cascade of phosphorylation events. In variant T207A, the absence of T207 phosphorylation impairs PLK1 docking explaining the reduction of binding to PLK1 and the global loss of phosphorylation by PLK1. S206C eliminates the serine residue at −1 position required for PLK1_PBD_ interaction resulting as well in a strong reduction of BRCA2 binding.

### Impairing T207 phosphorylation prolongs mitosis

PLK1 is a master regulator of mitosis ^17^. To find out whether the interaction between BRCA2 and PLK1 is involved in the control of mitotic progression we examined the functional impact of the variants that reduce PLK1 phosphorylation at T207 (S206C and T207A) in the context of the full-length BRCA2 protein in cells. For this purpose, we generated stable cell lines expressing the BRCA2 cDNA coding for either the GFPMBP-BRCA2 WT or the variants to complement DLD1 BRCA2 deficient human cells (hereafter BRCA2^-/-^). In this cell line, both alleles of BRCA2 contain a deletion in exon 11 causing a premature stop codon after BRC5 and cytoplasmic localization of a truncated form of the protein ^32^. We selected two stable clones of each variant that show similar protein levels as the BRCA2 WT complemented cells (clone C1, hereafter BRCA2 WT) by western blot (Supplementary Fig. 4a). We then tested the interaction of full-length BRCA2 with PLK1 in these stable clones by GFP pull-down assay. As expected, PLK1 readily co-purified with full-length BRCA2 WT from mitotic cells. Importantly, in cells expressing the variants S206C and T207A, the level of co-purified PLK1 was greatly reduced (Fig. 4a, 4b), confirming the results obtained with the overexpressed BRCA2_1-250_ fragments (Fig. 3a) now in the context of cells stably expressing the full-length BRCA2 protein. Thus, BRCA2 interaction with endogenous PLK1 is impaired in cells bearing variants S206C and T207A.

**Figure 4.**
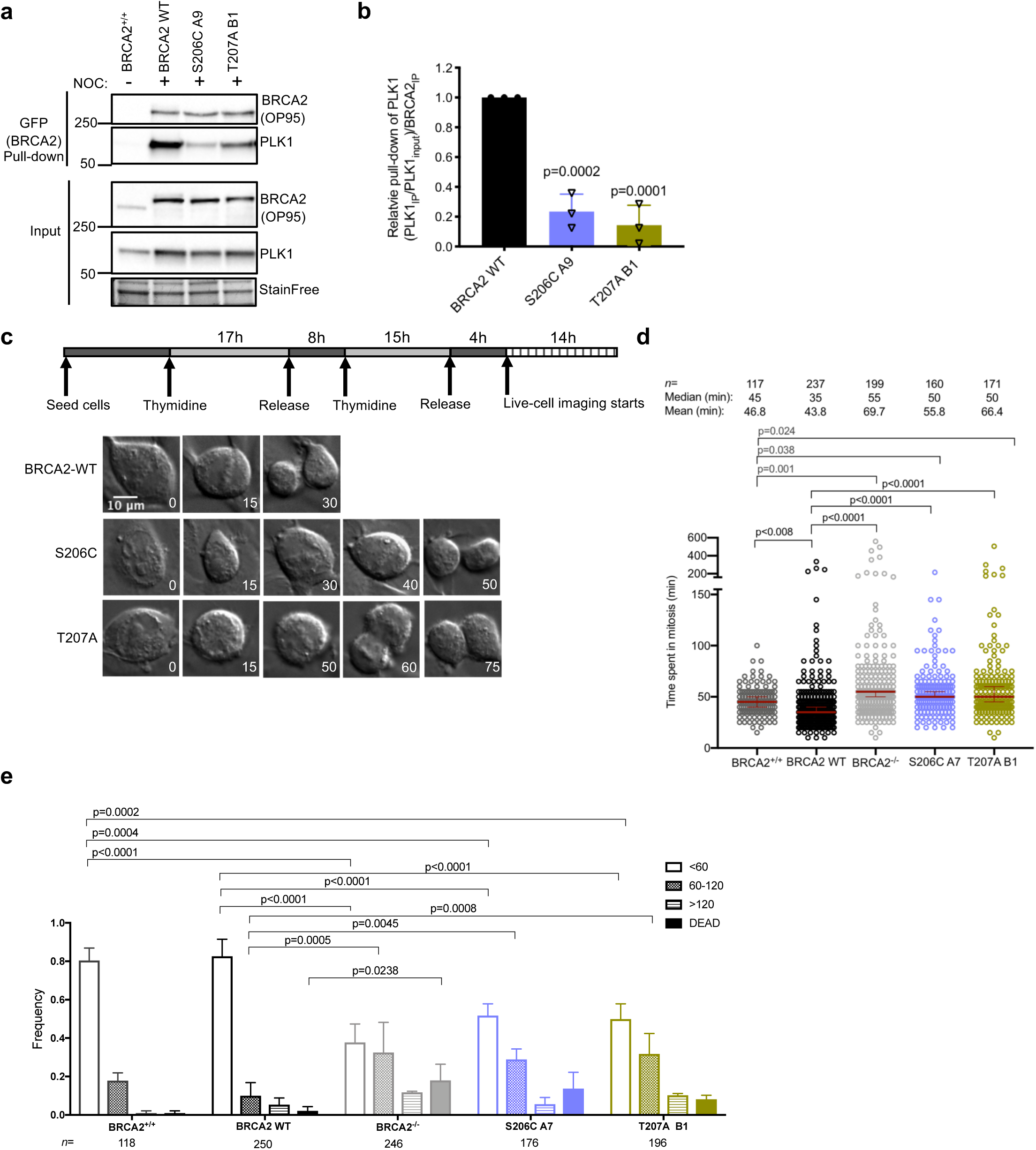
Cells bearing BRCA2 variants S206C and T207A prolong mitosis. **(a)** Interaction of full-length BRCA2 and PLK1 in stable clones of BRCA2 deficient DLD1 cells (BRCA2^-/-^) expressing GFPMBP-BRCA2 WT (BRCA2 WT) or the variants S206C or T207A. EGFPMBP-BRCA2 was immunoprecipitated from whole cell lysates of nocadozole treated cells using GFP-trap beads. Immuno-complexes were resolved on 4-15% SDS-PAGE followed by western blotting using anti-BRCA2 and -PLK1 antibodies. Unsynchronized DLD1 cells with endogenous BRCA2 (BRCA2^+/+^) were used as control for the immunoprecipitation and StainFree images of the gels before transfer were used as loading control for the input (cropped image is shown). **(b)** Quantification of co-immunoprecipitated PLK1 with EGFP-MBP-BRCA2 in (a) relative to the PLK1 protein levels in the input and the amount of immunoprecipitated (IP) EGFP-MBP-BRCA2 ((PLK1_IP_ /PLK1_input_)/EGFP-MBP-BRCA2_IP_) Results are presented as the fold change compared to the GFPMBP-BRCA2 WT clone. The data represents the mean ± SD of three independent experiments. Statistical significance of the difference between BRCA2 WT and the variants was calculated one-way ANOVA test with Dunnett’s multiple comparisons test. **(c)** Top: Scheme of the double thymidine block procedure used to synchronize DLD1 cells expressing endogenous BRCA2 (BRCA2^+/+^) or stable clones of DLD1 BRCA2 deficient cells (BRCA2^-/-^) expressing GFPMBP-BRCA2 WT or VUS as indicated, for live-cell imaging of mitotic progression. Bottom: Representative still images of the live-cell imaging at different time points. The numbers in each image indicate the time in minutes after nuclear envelope break down (NEBD). Scale bar represents 10 µm. **(d-e)** Quantification of the time the cells spent in mitosis from nuclear envelope break down to mitotic exit monitored by time-lapse microscopy. (d) Scatter dot plot showing the quantification of the live-time imaging in (c). The median time (min) the dividing cells spent in mitosis is indicated by a red line in the plot (median with 95% CI). Each dot represents a single cell, *n* indicates the total number of cells counted from two to four independent experiments (BRCA2^+/+^ (n=2), WT C1 (n=5), BRCA2^-/-^ (n=4), S206C A7 (n=3), T207A B1 (n=3)), with 45 to 60 cells counted per experiment. For statistical comparison of the differences between the samples we applied a Kruskal-Wallis test followed by Dunn’s multiple comparison test, the p-values show significant differences. (**e**) Frequency distribution of the time spent in mitosis, including cells that fail to divide (DEAD). The error bars represent SD from at least two-four independent experiments (as in d). Statistical significance of the difference in (d and e) was calculated with one-and two-way ANOVA test respectively, with Tukey’s multiple comparisons test. The p-values show significant differences.

Also, the binding of BRCA2 to PLK1 in cells stably expressing full-length BRCA2 WT was not reduced by incubating the cells with PLKi (BTO) (Supplementary Fig. 4b), consistently with results obtained *in vitro* (Fig. 3c). Altogether, these data suggest that another binding site primed by a different kinase (presumably T77 phosphorylated by CDK1^33^) contributes to BRCA2 binding to PLK1. Consistent with this idea, the binding of overexpressed 2xMBP-BRCA2_1-250_ to the endogenous PLK1 in U2OS cells was completely abolished in the presence of the CDK inhibitor (RO3306) (Supplementary Fig. 4c, lane 4 compared to lane 2).

Having confirmed that T207 was a docking site for PLK1 in cells, we next examined the impact of BRCA2 variants on mitosis. Therefore, we monitored the time taken for individual cells from mitotic entry (defined as nuclear envelope break down) to mitotic exit using live cell imaging. Cells expressing the endogenous BRCA2 (hereafter BRCA2^+/+^) and the BRCA2 WT cells showed similar kinetics, they completed mitosis, on average, in 47 and 44 min, respectively (Fig. 4c, d) and the majority of the cells (80% for BRCA2^+/+^ and 82% for BRCA2 WT) completed mitosis within 60 min (Fig. 4e). In contrast, cells expressing variants S206C and T207A augmented the time spent in mitosis (average time of 56 and 66 min, respectively, Fig. 4d). This trend was also observed in the frequency of cells dividing within 60 min (∼49-51%), compared to 82% in BRCA2 WT cells (Fig. 4e). Representative videos of the still images shown in Fig. 4c are included in Supplementary movies 1-3.

Taken together, cells expressing variants S206C and T207A display a significant delay in mitotic progression compared to BRCA2 WT expressing cells.

### Docking of PLK1 at T207 of BRCA2 favours the formation of a complex with PP2A and phosphorylated BUBR1 that facilitates chromosome alignment

BRCA2 forms a complex with BUBR1^4, 5^. BUBR1 facilitates kinetochore-microtubule attachments via its interaction with the phosphatase PP2A. Phosphorylation of BUBR1 by PLK1 at the KARD motif comprising the tension sensitive sites S676 and T680 promotes interaction with PP2A^20^. A defect in the phosphorylation of BUBR1 weakens its interaction with PP2A leading to mitotic delay^20, 34^. The mitotic delay phenotype we observed in BRCA2 mutated cell lines led us to ask whether BRCA2 and PLK1 formed a tetrameric complex with pBUBR1 and PP2A. Using a GFP pull-down to capture GFP-MBP-BRCA2 from mitotic BRCA2 WT cells, we observed that PLK1, pT680-BUBR1 and PP2A (detected with an antibody against the catalytic subunit of PP2A, PP2AC) were pull-down together with GFP-MBP-BRCA2, indicating the formation of a tetrameric complex (Fig. 5a). As described for pBUBR1^19, 20^, PLK1^34^ and PP2A^22^, we found BRCA2 at the kinetochore in mitotic cells (Supplementary Fig. 5a) as previously reported^4^ supporting the idea that this complex takes place at the kinetochore.

**Figure 5.**
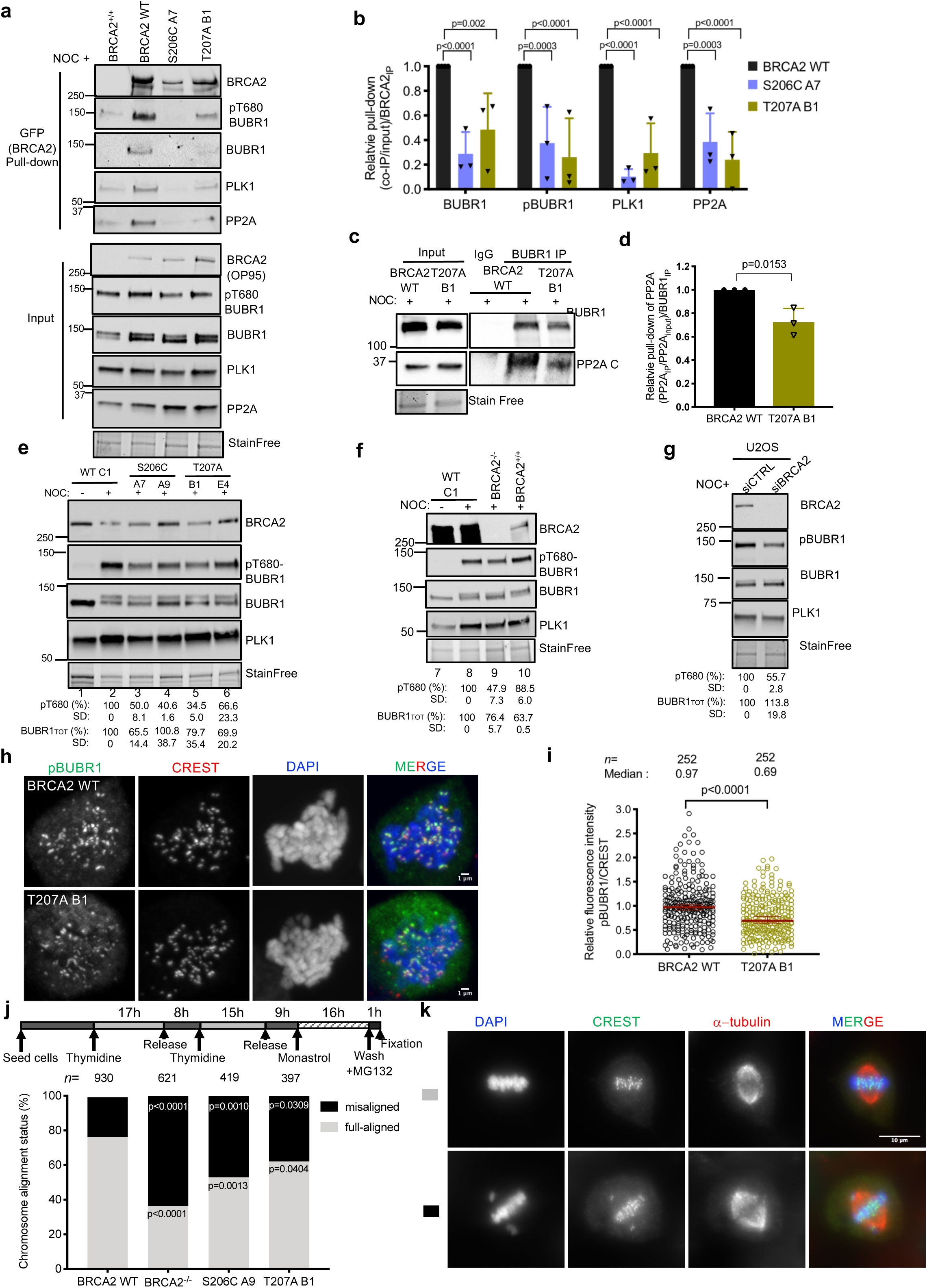
Cells expressing BRCA2 variants that alter PLK1 phosphorylation display reduced protein levels of phosphorylated BUBR1 and its interaction with PP2A resulting in chromosomes misalignment. **(a)** Co-immunoprecipitation of the protein complex BRCA2-BUBR1/pBUBR1-PP2A-PLK1 from mitotic cell extracts of BRCA2 WT cells or cells expressing the variant S206C and T207A using GFP-trap beads. The immuno-complexes were resolved on 4-15% SDS-PAGE followed by western blotting, the interactions were revealed by anti-BRCA2, -BUBR1, - pT680-BUBR1, -PLK1 and -PP2AC (PP2A catalytic subunit) antibodies. Mitotic DLD1 cells with endogenous BRCA2 (BRCA2^+/+^) were used as control for the immunoprecipitation. **(b)** Quantification of co-immunoprecipitated BUBR1, pBUBR1, PLK1 and PP2A with EGFPMBP-BRCA2 in (a), relative to the input levels of each protein and the amount of immunoprecipitated EGFP-MBP-BRCA2. Results are presented as the fold change compared to the BRCA2 WT clone. The data represents the mean ± SD of three independent experiments. Statistical significance of the difference was calculated with two-way ANOVA test with Dunnett’s multiple comparisons test, the p-values show the significant difference. **(c)** Co-immunoprecipitation of endogenous PP2AC with endogenous BUBR1 from mitotic cell extracts of BRCA2 WT cells or cells expressing the variant T207A using mouse anti-BUBR1 antibody. Mouse IgG was used as control for the BUBR1 immunoprecipitation. The immuno-complexes were resolved on 4-15% SDS-PAGE followed by western blotting, the interactions were revealed by rabbit anti-BUBR1 and anti-mouse PP2AC antibodies. **(d)** Quantification of co-immunoprecipitated PP2A C in (c), relative to the input levels and the amount of immunoprecipitated BUBR1. Results are presented as the fold change compared to the BRCA2 WT clone. The data represents the mean ± SD of three independent experiments. Statistical significance of the difference was calculated with unpaired two-tailed t-test, the p-values show the significant difference. **(e-g)** Western blots showing the expression levels of endogenous BUBR1 and pT680-BUBR1 in nocodazole treated stable clones of DLD1 BRCA2 deficient cells (BRCA2^-/-^) expressing GFPMBP-BRCA2 WT (BRCA2 WT) or the variants, as indicated (e) or in nocodazole treated DLD1 BRCA2 deficient cells (BRCA2^-/-^) or DLD1 cells expressing endogenous BRCA2 (BRCA2^+/+^) (f). **(g)** Western blot showing the expression levels of endogenous BUBR1 and pT680-BUBR1 in U2OS after down-regulation of endogenous BRCA2 using siRNA. (e-g) The mean BUBR1_TOT_ and pBUBR1 signal relative to the stain free signal is shown for the nocadozole treated samples below the blots, results are presented as percentage compared to BRCA2 WT clone. The data represents the mean ± SD of three (e) and two (g and g) independent experiments. The protein levels of PLK1 in (a, e-g) are shown as a G2/M marker. **(h)** Representative images of the localization of pT680-BUBR1 in nocodazole-arrested DLD1 BRCA2^-/-^ cells stably expressing GFP-MBP-BRCA2 WT or the variant T207A as indicated. CREST is used as centromere marker and DNA is counterstained with DAPI. Scale bar represents 1 µm. **(i)** Quantification of the co-localization of pT680-BUBR1 and CREST in (h). The data represents the intensity ratio (pT680-BUBR1:CREST) relative to the mean ratio of pT680-BUBR1:CREST for the GFP-MBP-BRCA2 WT calculated from a total of 252 pairs of chromosomes analysed from two independent experiments (6 pairs of chromosomes/cell from 21 cells). The red line in the plot indicates the median (95% CI) ratio, each dot represents a pair of chromosomes. For statistical comparison of the differences between the samples we applied a Mann-Whitney test, the p-values show significant differences. **(j)** Top: Scheme of the double thymidine block procedure used to synchronize the DLD1 cells for the analysis of chromosome alignment. Bottom: Quantification of misaligned chromosomes outside the metaphase plate in DLD1 BRCA2 deficient cells (BRCA2^-/-^) an BRCA2^-/-^ cells stably expressing BRCA2 WT or the S206C and T207A variants. *n* indicates the total number of cells counted for each clone from two (BRCA2^-/-^, S206C and T207A) and four (BRCA2 WT) independent experiments. Statistical significance of the difference was calculated with unpaired two-way ANOVA test with Tukey’s multiple comparisons test, the p-values show the significant difference. **(k)** Representative images of the type of chromosome alignment observed in cells quantified in (j), scale bar represents 10 µm.

Importantly, cells expressing the variants S206C or T207A showed a strong reduction in the interaction of BRCA2 with PLK1, PP2A, BUBR1 and pT680-BUBR1 in the context of the tetrameric complex (Fig. 5a, b). Moreover, the overall levels of BUBR1 and pBUBR1 were also reduced in cells bearing S206C and T207A variants compared to the WT cells, as detected by specific antibodies against BUBR1, pT680-BUBR1 (Fig. 5e-i) and pS676-BUBR1 (Supplementary Fig. 5d), and this was also the case in BRCA2 deficient cells (DLD1 BRCA2^-/-^ cells) or U20S cells depleted of BRCA2 by siRNA (Fig. 5f, g). Furthermore, we observed an overall reduction in the levels of pBUBR1 at the kinetochore (Fig. 5h, i) in cells expressing T207A compared to WT cells. Consistently, when we immunoprecipitated BUBR1 from mitotic cells and detected the levels of co-immunoprecipitated PP2A (PP2AC antibody), we observed that, although PP2A was readily copurified with BUBR1 in the BRCA2 WT cells, expressing BRCA2 variant T207A reduced the levels of PP2A by ∼30% (Fig. 5c, 5d) suggesting that BRCA2 facilitates the formation of a complex between BUBR1-and PP2A.

The association of phosphorylated BUBR1 with PP2A is required for the formation of stable kinetochore-microtubule attachments ^19, 20^, a defect in this interaction resulting in chromosome misalignment. Therefore, we next examined whether cells expressing the BRCA2 variants S206C and T207A altered chromosome alignment. Following thymidine synchronization, the cells were treated with the Eg5 inhibitor Monastrol (100 μM) for 14h followed by Monastrol washout and release for 1h in normal media supplemented with the proteasome inhibitor MG132 to avoid exit from mitosis ^19^. Chromosome alignment was then analysed by immunofluorescence. Importantly, the analysis of cells expressing S206C and T207A variants showed high frequency of faulty chromosome congression compared to the BRCA2 WT clone (47% in S206C and 38% in T207 versus 24% in the BRCA2 WT clone), which was exacerbated in BRCA2^-/-^ cells (63%) (Fig. 5j, k), as detected by signals of the centromere marker (CREST) outside the metaphase plate (Fig. 5k).

Mutations in the PLK1-targeted sites of BUBR1 lead to chromosome misalignment, a phenotype that can be rescued by the phosphomimic mutant BUBR1-3D (S670D, S676D, T680D) ^20^. To better understand the mechanism behind the phenotype observed in our clones, we overexpressed RFP-BUBR1-3D to test whether this form of BUBR1 could restore the misalignment phenotype observed in cells bearing T207A. Surprisingly, the cells overexpressing RFP-BUBR1-3D presented a similar misalignment phenotype, ∼38% of exhibited misaligned chromosomes compared to 33% for cells bearing T207A (Fig. 6a, b). Importantly, BUBR1-3D, which has a weaker affinity for PP2A than BUBR1 phosphorylated by PLK1 *in vitro*^35^, but is sufficient to restore PP2A binding in BUBR1 deficient cells ^20^, could not rescue the interaction of BUBR1 with PP2A in cells bearing BRCA2 S206C variant (Fig. 6c).

**Figure 6.**
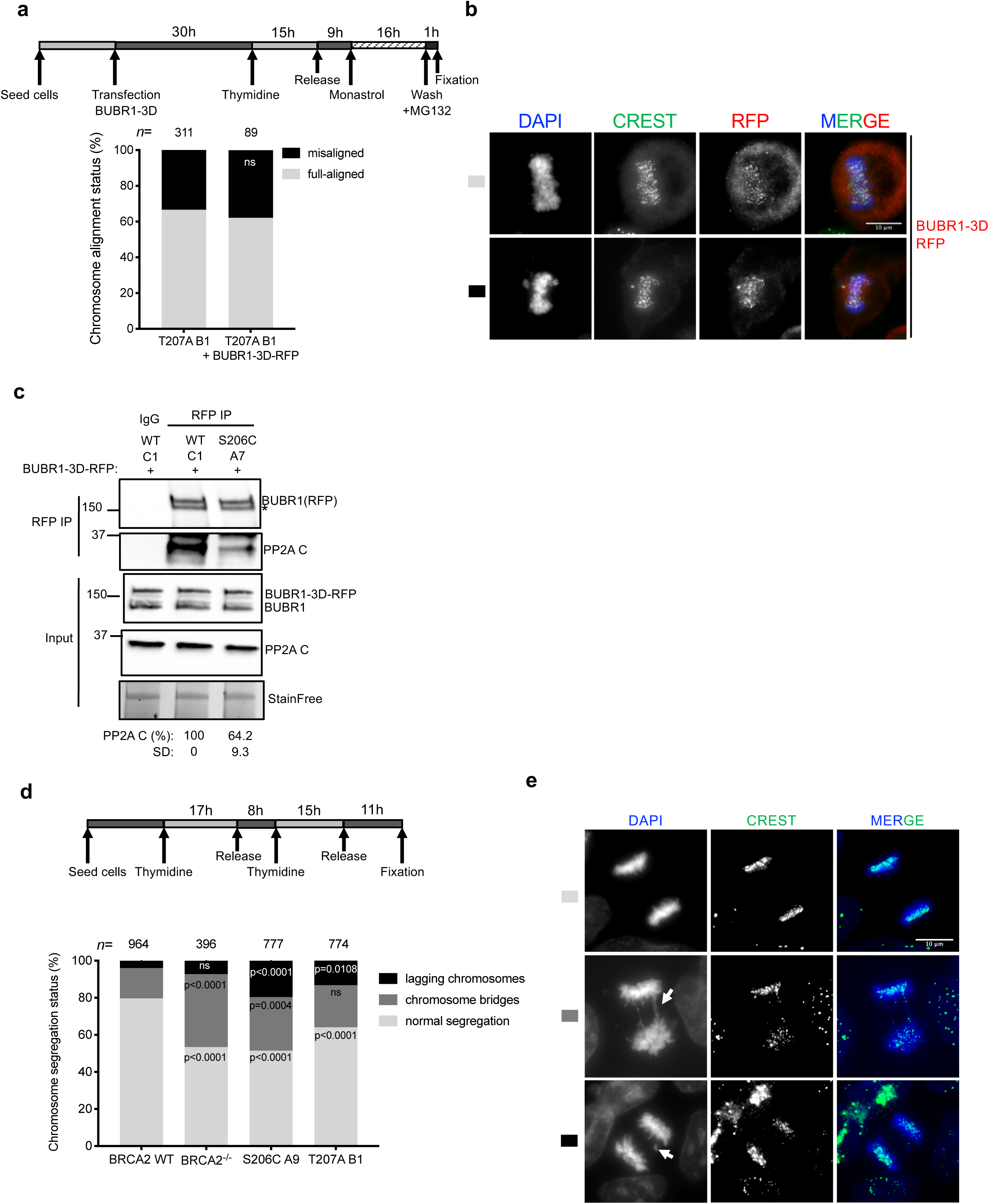
Cells expressing BRCA2 variants S206C and T207A display aberrant chromosome segregation. **(a)** Top: Scheme of the synchronization procedure for the analysis of chromosome alignment in the T207A cells transient overexpressing the BUBR1-3D mutant (S670D, S676D and T680D). Bottom: Quantification of misaligned chromosomes outside the metaphase plate in DLD1 BRCA2^-/-^ T207A stable clones transient overexpressing the 3xFLAG-BUBR1-3D-RFP mutant (S670D, S676D and T680D). *n* indicates the total number of cells counted for each clone from two independent experiments. Statistical significance of the difference was calculated with unpaired two-tailed t-test, ns (non-significant)). **(b)** Representative images of the type of chromosome alignment observed in cells overexpressing 3xFLAG-BUBR1-3D-RFP quantified in (a); scale bar represents 10 µm. (**c)** Co-immunoprecipitation of endogenous PP2AC (PP2A catalytic subunit) with transient overexpressed 3xFLAG-BUBR1-3D-RFP mutant (S670D, S676D and T680D) from mitotic cell extracts of BRCA2 WT cells or cells expressing the S206C variant using rabbit anti-RFP antibody. Rabbit IgG was used as control for the BUBR1 immunoprecipitation. The immuno-complexes were resolved on 4-15% SDS-PAGE followed by western blotting, the interactions were revealed by mouse anti-BUBR1 and PP2AC antibodies. The amount of PP2A co-immunoprecipitated with BUBR1-3D-RFP relative to the input levels of PP2A and the amount of immunoprecipitated BUBR1-3D-RFP is presented below the blot as mean ± SD from two independent experiments. The data is presented relative to the non-treated BRCA2 WT. **(d)** Top: Scheme of the double thymidine block procedure used to synchronize DLD1 BRCA2^-/-^ stable clones for analysis of aberrant chromosome segregation. Bottom: Quantification of cells with aberrant chromosomes segregation in BRCA2^-/-^ cells and in the clones stably expressing BRCA2 WT, S206C and T207A, as indicated. Statistical significance of the difference in (d) was calculated with two-way ANOVA test with Tukey’s multiple comparisons test (the p-values show the significant differences compared to WT; ns (non-significant)). *n* in (d) indicates the total number of cells counted for each clone from two (BRCA2^-/-^, S206C and T207A) and four (BRCA2 WT) independent experiments. **(e)** Representative images of the type of aberrant chromosome segregation observed in the cells quantified in (d), CREST antibody is used as marker of centromere; nuclei are revealed with DAPI counterstaining. Scale bar represents 10 µm.

Our results strongly suggest that docking of PLK1 onto BRCA2 T207 facilitates the formation of a complex between phosphorylated BUBR1 and PP2A at the kinetochore. This is not due to an increased localization of PLK1 at the kinetochore, as the levels of PLK1 at the kinetochores remained unchanged in BRCA2 WT and variant (Supplementary Fig. 6a, b), nor to an increased interaction of PLK1 with BUBR1 (Supplementary Fig. 6c, d).

### The variants that reduce PLK1 phosphorylation of BRCA2 display strong defects in chromosome segregation and aneuploidy

Unresolved chromosome misalignment as observed in cells altering BRCA2 phosphorylation by PLK1 is expected to drive chromosome missegregation. To find out if this was the case in cells expressing BRCA2 variants S206C and T207A, we examined chromosome segregation by immunofluorescence in cells synchronized by double-thymidine block and released for 15h to enrich the cell population at anaphase/telophase stage. BRCA2^-/-^ cells displayed, as expected, an increased proportion of chromosome bridges (39% vs 16% in cells expressing BRCA2 WT), whereas the fraction of lagging chromosomes was only mildly increased (7% vs 4% in BRCA2 WT). The percentage of chromosome bridges in cells expressing S206C and T207A was moderately increased (23% and 29%, respectively, compared to 16% in BRCA2 WT). However, the biggest difference was observed in the percentage of lagging chromosomes increasing between 3-and 5-fold in the cells bearing the variants compared to the BRCA2 WT cells (Fig. 6d, 6e).

Erroneous chromosome segregation generates aneuploid cells during cell division ^36^. Given the strong chromosome segregation defects observed in cells expressing S206C and T207A we next analysed the number of chromosomes in these cells. Total chromosome counts carried out on metaphase spreads revealed that 37.1% of BRCA2 WT cells exhibited aneuploidy with chromosome losses or gains. In the case of S206C and T207A, this number was elevated to 52.2% and 61.8% of the cells, respectively (Fig. 7a). An example of the images analyzed can be found in Fig. 7b. As the number of chromosomes was difficult to assess for cells with high content of chromosome gains we arbitrarily discarded cells that contained more than 65 chromosomes. Thus, tetraploid cells were not included in this measurement. Therefore, as a complementary test, we determined the frequency of tetraploid cells by assessing the incorporation of BrdU and measuring the frequency of S-phase cells with >4N DNA content (Fig. 7c). Quantification of these data showed that similar to the BRCA2 WT complemented cells, the frequency of tetraploidy in cells bearing the variants is <1% of the total population (Fig. 7d), while the frequency of BrdU positive cells was equivalent between the BRCA2 WT and the VUS expressing cells (Supplementary Fig. 7).

**Figure 7.**
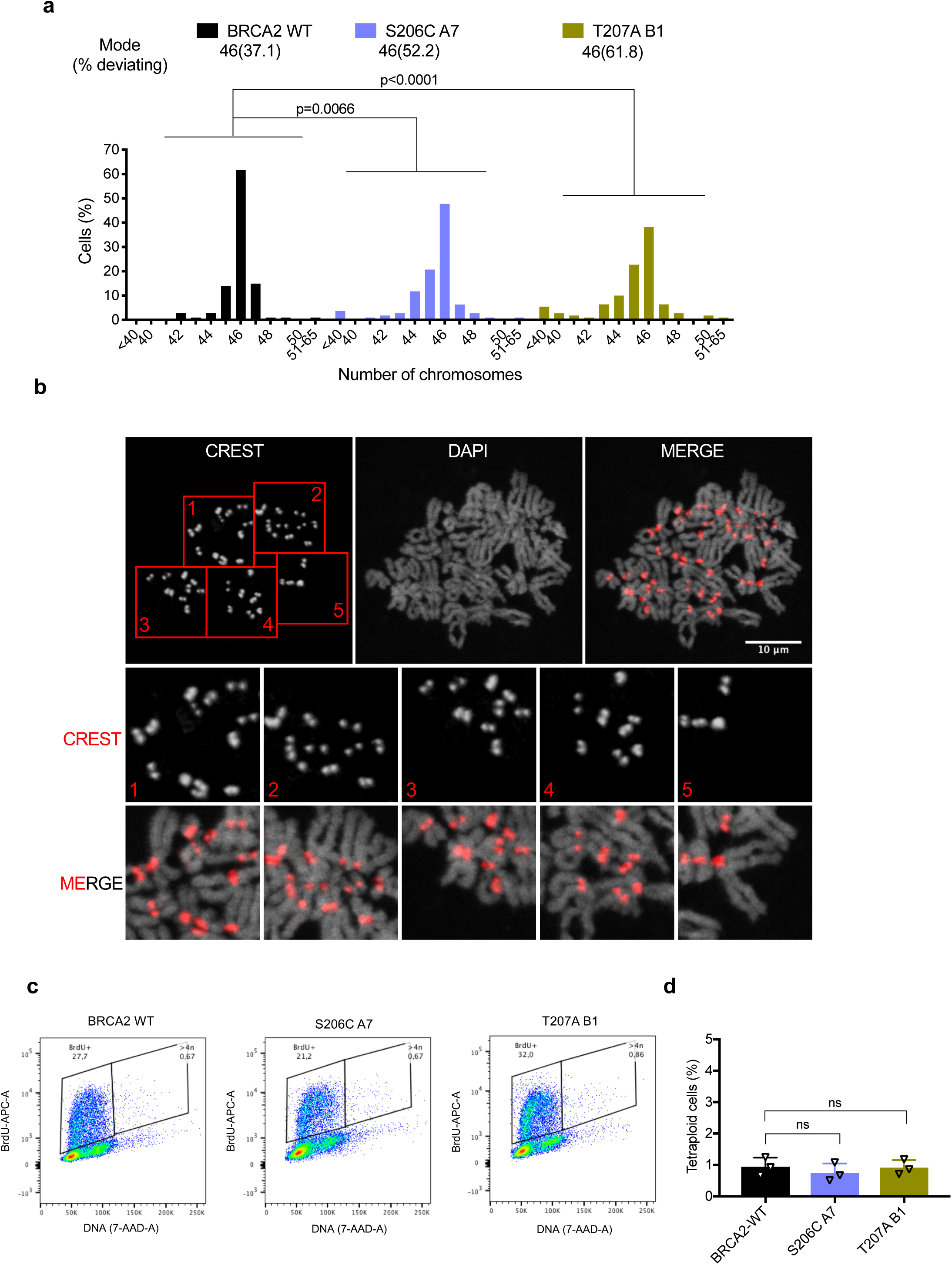
Cells expressing BRCA2 variants S206C and T207A exhibit aneuploidy. **(a)** Distribution of the number of chromosomes observed in metaphase spreads of stable clones expressing BRCA2 WT, S206C A7 or T207 B1 from two independent experiments (total number of cells counted; BRCA2 WT (n=105), S206C A7 (n=111) and T207A B1 (n=110)); modal number of chromosomes and percentage deviating from the mode are shown at the top; Kruskal-Wallis test followed by Dunn’s multiple comparison test was used for statistical comparison of the differences between the samples, the p-values show the significant differences.). The cell passage was between 5 and 9 (BRCA2 WT (p.6 and p.9), S206C A7 (p5 and p9) and T207A B1 (p.6 and p.9). **(b)** Representative image of metaphase spreads of the DLD1 BRCA2 deficient stable cells bearing the S206C BRCA2 variant stained with CREST and counterstained with DAPI. In this example, the cell contains 45 chromosomes. **(c-d)** Analysis of S-phase tetraploid cells in DLD1 BRCA2 deficient cells expressing BRCA2 WT or the VUS S206C and T207A measured by flow cytometry after 20 minutes of BrdU incorporation. **(c)** Representative flow cytometry plots of cells stained with anti-BrdU-APC antibodies and 7-AAD (DNA). **(d).** Frequency of S-phase tetraploid cells in stable clones expressing BRCA2 WT or the VUS S206C and T207A. The data represents the mean ± SD of three independent experiments (cell passage: 6-10). Statistical significance of the difference was calculated with one-way ANOVA test with Tukey’s multiple comparisons test (the p-values show the difference compared to WT, ns: non-significant).

Together, these results indicate that, in addition to the severe chromosome misalignment phenotype, cells expressing S206C and T207A display high frequency of chromosome missegregation, including a strong induction of lagging chromosomes and a mild increase in chromosome bridges. As a consequence, the incidence of aneuploidy, but not tetraploidy, is greatly exacerbated in these cells.

### The variants altering PLK1 phosphorylation of BRCA2 restore the hypersensitivity of BRCA2 deficient cells to DNA damage and PARP inhibition and are HR proficient

Since BRCA2 has a major role in DNA repair by HR, the prolonged mitosis observed in the VUS-expressing stable cell lines (Fig. 4) could result from checkpoint activation through unrepaired DNA. In addition, defects in chromosome segregation have been reported as a cause of DNA damage. Thus, to test the DNA repair efficiency of cells expressing variants S206C and T207A, we performed a clonogenic survival assay in the stable clones after treatment with mitomycin C (MMC), an inter-strand crosslinking agent to which BRCA2 deficient cells are highly sensitive ^37^. As expected, BRCA2 deficient cells (BRCA2^-/-^) showed hypersensitivity to MMC treatment whereas BRCA2 WT cells complemented this phenotype almost to the same survival levels as the cells expressing the endogenous BRCA2 (BRCA2^+/+^). The stable clones expressing variants S206C and T207A also complemented the hypersensitive phenotype of BRCA2^-/-^ cells, although there was a mild reduction in the survival levels compared to the BRCA2 WT cells (Fig. 8a), consistent with the slightly increased levels of chromosome bridges (Fig. 6d). These results suggest that the delay in mitosis is not a consequence of checkpoint activation via unrepaired DNA. Cells expressing VUS S206C and T207A showed a growth defect manifested in a reduced number of colonies observed in unchallenged conditions (Supplementary Fig. 8a), which is consistent with the mitotic phenotype observed (Fig. 4-6). To exclude a possible bias arising from the different ability to form colonies of these cells we also tested the sensitivity to MMC of the cell population by MTT assay. As shown in Fig. 8b, cells expressing S206C and T207A showed similar relative viability compared to BRCA2 WT complemented cells or the cells expressing endogenous BRCA2 (BRCA2^+/+^), confirming our results. To additionally address the DNA repair capacity of these cells, we tested their viability upon treatment with the poly (ADP-ribose) polymerase (PARP) inhibitor Olaparib. PARP1 is an enzyme required for the sensing of DNA single strand breaks (SSBs) and double strand breaks (DSBs) that becomes essential in the absence of a functional HR pathway ^38^ and therefore is used as a surrogate of HR proficiency. In fact, PARP1 inhibitors, in particular Olaparib, is currently used in the clinic to treat breast and ovarian cancer patients carrying germline mutations in BRCA1/2. In our settings, the relative viability of BRCA2^-/-^ cells was 45% upon 4-day treatment with the highest Olaparib concentration tested (5 µM); in contrast, 74% of BRCA2 WT complemented cells remained viable. Similarly, cells expressing S206C or T207A survived the treatment equally well as the cells expressing BRCA2 WT, the percentage of viable cells at 5 µM treatment ranging from 70 to 83 % depending on the clone (Fig. 8c) indicating that the mutated cells can rescue the phenotype of BRCA2^-/-^ cells.

**Figure 8.**
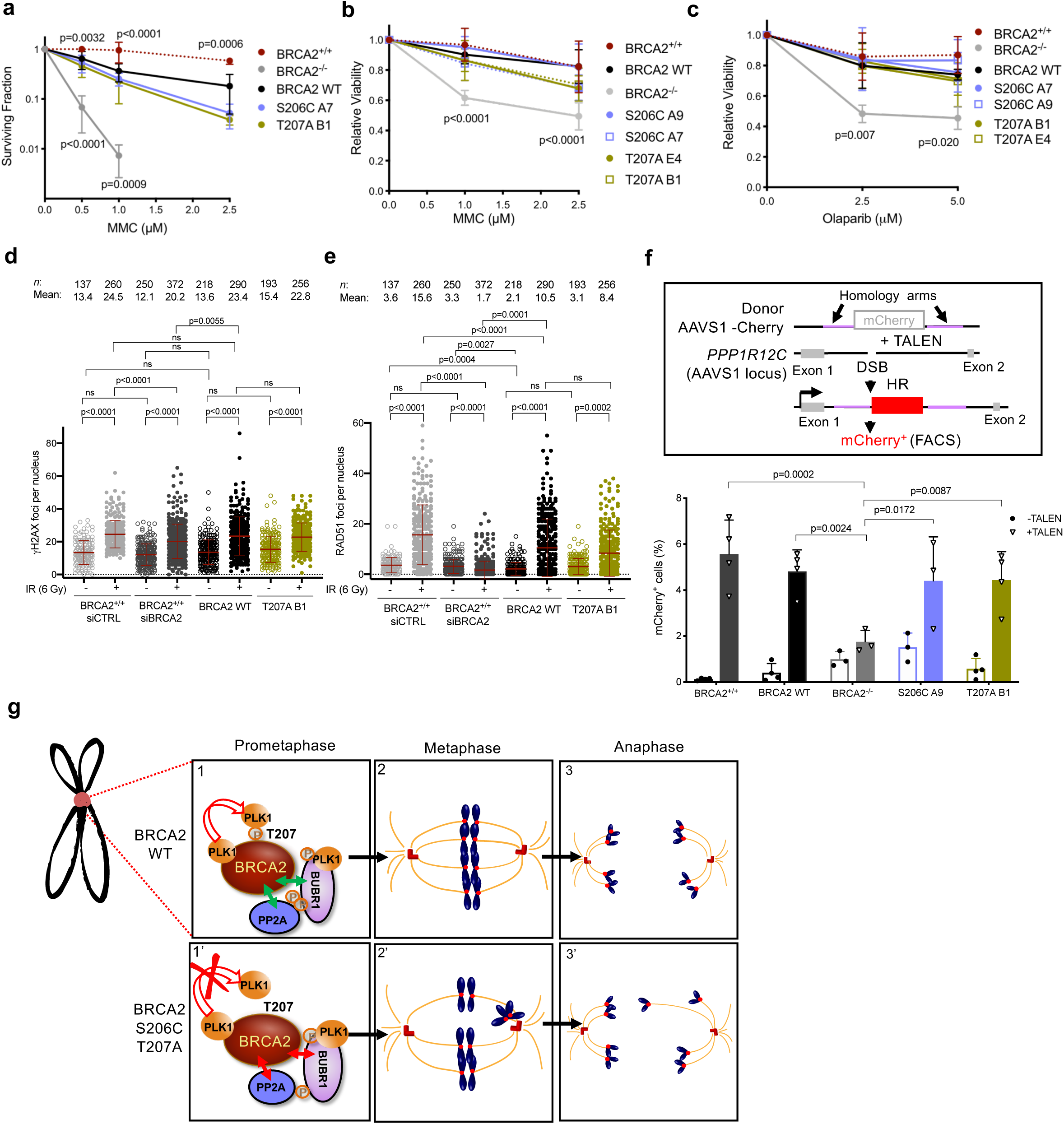
The DNA repair proficiency is not affected in cells bearing BRCA2 variants S206C and T207A. **(a)** Quantification of the surviving fraction of DLD1 cells expressing endogenous BRCA2 (BRCA2^+/+^) or stable clones of BRCA2 deficient DLD1 cells (BRCA2^-/-^) expressing BRCA2 WT or the variants S206C or T207A assessed by clonogenic survival upon exposure to MMC at concentrations: 0, 0.5, 1.0 and 2.5 µM. Data are represented as mean ± SD: BRCA2^+/+^ (red) (n=3), BRCA2^-/-^ (gray) (n=6), WT C1 (black) (n=6), S206C A7 (blue) (n=3), T207A B1 (green) (n=4). **(b-c)** Quantification of the relative cell viability monitored by MTT assay upon treatment with increasing doses of MMC (b) or the PARP inhibitor Olaparib (c), as indicated. The data represents the mean ± SD of four (b) and three (c) independent experiments. Statistical significance of the difference in (a-c) was calculated with two-way ANOVA test with Tukey’s multiple comparisons test (the p-values show the significant differences compared to the BRCA2 WT clone). **(d-e)** Quantification of the number of γH2AX foci (d) or RAD51 foci per nucleus (e) two hours after 6 Gy of γ-irradiation (+IR) versus non-irradiated conditions (-IR), in DLD1 BRCA2^+/+^ cells depleted of BRCA2 (siBRCA2) or control cells (siCTRL) or BRCA2 deficient DLD1 cells (BRCA2^-/-^) expressing BRCA2 WT or the variant T207A. *n* indicates the total number of cells counted from two independent experiments. For statistical comparison of the differences between the samples we applied a Kruskal-Wallis test followed by Dunn’s multiple comparison test, the p-values show significant differences. The red line in the plot indicates the mean ± SD, each dot represents a single focus. **(f)** Top: Scheme of the DSB-mediated gene targeting HR assay. Bottom: Frequency of mCherry positive cells in cells transfected with the promoter-less donor plasmid (AAVS1-2A-mCherry) without (-TALEN) or with (+TALEN) nucleases. Statistical significance of the difference in (f) was calculated with two-way ANOVA test with Tukey’s multiple comparisons test, the p-values show the significant differences. **(g)** Model for the role of PLK1 phosphorylation of BRCA2 T207A by PLK1 in mitosis (see text for details). In panel 1 and 1’ the two-sided arrows represent a complex between the two proteins as indicated either favored by BRCA2 WT (green) or impaired in the BRCA2 mutated form (red); in panels 2, 2’, 3 and 3’ blue blobs represent chromosomes, red circles represent the kinetochores, red cylinders represent the centrioles and orange lanes represent the spindle microtubules.

To determine directly the levels of spontaneous DNA damage in these cells and whether or not they can recruit the recombination protein RAD51 to sites of DNA damage, we measured both the number of nuclear foci of both the DSB marker γH2AX and RAD51 protein, in cells unchallenged (-IR) or 2h upon induction of DNA damage by ionizing radiation (6 Gy, (+IR)). Our results show that the number of γH2AX foci in unchallenged conditions, indicative of spontaneous DNA damage, is comparable in all cell lines including in cells transiently depleted of BRCA2 by siRNA (Fig. 8d and Supplementary Fig. 8b); this is probably due to the high genome instability intrinsic to these cancer cells. Similarly, the number of RAD51 foci in unchallenged conditions did not change significantly among samples (mean varying between 2-3 foci). In contrast, the number of RAD51 foci upon irradiation increased between 5-fold in DLD1 BRCA2^+/+^ cells and BRCA2 WT cells and 3-fold in T207A bearing cells whereas in cells depleted for BRCA2 the number of RAD51 foci decreased to 1 focus per nucleus, on average (Fig. 8e) whereas the number of γH2AX foci upon irradiation among cells remained the same (Fig. 8d). From these results, we conclude that the DNA repair foci are only mildly altered in cells expressing T207A. Representative images of these experiments are shown in Supplementary Fig. 8b.

A typical feature of replication stress is the appearance of micronuclei in daughter cells which generally contain DNA fragments. In contrast, the presence of micronuclei with centromeres is an indication of whole chromosome or chromatid, suggesting an event arising from lagging chromosomes. We measured the number of micronuclei with and without centromeres in our cells. BRCA2 deficient cells displayed an overall increased number of micronuclei compared to BRCA2 WT cells. In contrast, cells bearing S206C or T207A variant did not exhibit an increase in micronuclei either with or without centromeres (Supplementary Fig. 8d, e) excluding strong replication stress-induced DNA damage in these cells.

Finally, to directly assess the HR proficiency of these cells, we performed a cell-based HR assay by DSB-mediated gene targeting at a specific locus (AAVS1 site) within the endogenous PPP1R12C gene using a site-specific transcription-activator like effector nuclease (TALEN) and a promoter-less mCherry donor flanked by homology sequence to the targeted locus ^39, 40^. DSB-meditated gene targeting results in mCherry expression from the endogenous PPP1R12C promoter (Fig. 8f) which can be measured by flow cytometry (Supplementary Fig. 9). Using this system, BRCA2^+/+^ and BRCA2 WT complemented cells show ∼ 7% of mCherry positive TALEN-transfected cells (mean of 5.6% for BRCA2^+/+^ and 4.9% for WT) whereas the BRCA2 deficiency (BRCA2^-/-^) led to a reduction to ∼2% of mCherry expressing cells, as expected. Importantly, TALEN-transfected cells expressing BRCA2 variants S206C and T207A showed no significant difference with the BRCA2 WT complemented cells indicating an intact HR activity.

In summary, these results indicate that the role of BRCA2 in conjunction with PLK1 in mitosis is likely independent of the HR function of BRCA2 as the variants S206C and T207A affecting PLK1 phosphorylation of BRCA2 are only mildly sensitive to DNA damage, do not show an increased number of micronuclei, are able to recruit RAD51 to DNA damage sites (as shown for T207A) and are efficient at DSB-mediated gene targeting.

## Discussion

Our results demonstrate that residues S193 and T207 of BRCA2 can be efficiently phosphorylated by PLK1 (Fig. 2), thus extending the consensus sequence for phosphorylation by this kinase: position 205 is a serine and not a negatively charged residue, as generally observed at PLK1 phosphorylation sites^24^. Moreover, we reveal that pT207 constitutes a *bona fide* docking site for PLK1_PBD_ (Fig. 3h-l). Accordingly, *BRCA2* missense variants of unknown clinical significance reducing the phosphorylation status of T207 (T207A, S206C) result in a decreased in BRCA2-PLK1 interaction (Fig. 3a-k, 4a). The phenotype of cells expressing these two breast cancer variants in a BRCA2 deficient background was characterized, in order to investigate in detail the possible role of BRCA2 phosphorylation at T207 by PLK1 in the control of mitosis. Unexpectedly, we found that the cells expressing S206C and T207A display defective chromosome congression to the metaphase plate (Fig. 5j, k), causing a substantial delay in mitosis progression (Fig. 4c-e).

Proper kinetochore-microtubule attachments require the interaction of BUBR1 with the phosphatase PP2A-B56α to balance Aurora B kinase activity ^20, 22^. This interaction is mediated through the phosphorylation of the KARD motif of BUBR1 by PLK1. BRCA2 does not alter PLK1 interaction with BUBR1 (Supplementary Fig. 6c, d). However, we found that BRCA2 forms a tetrameric complex with PLK1-pBUBR1-PP2A, and that this complex is strongly reduced in cells bearing BRCA2 variants S206C and T207A (Figure 5a, 5b). Furthermore, cells bearing BRCA2 variants S206C and T207A show reduced overall levels of BUBR1 including pBUBR1(pT680) at the kinetochore (Fig. 5i). Importantly, the fact that in the BRCA2 mutated cells the interaction of PP2A with total BUBR1 is reduced (Fig. 5c, d) and that neither this interaction (Fig. 6c) nor the chromosome alignment defect can be rescued by BUBR1-3D overexpression (Fig. 6a, b) strongly suggests that PP2A needs to be in complex with PLK1-bound BRCA2 to bind BUBR1 and facilitate chromosome alignment. Cells bearing T207A and S206C variants display chromosome segregation errors including lagging chromosomes and chromosome bridges (Fig. 6d, e). Importantly, these accumulated errors ultimately lead to a broad spectrum of chromosome gains and losses (aneuploidy) compared to the wild type counterpart (Fig. 7a), but not to tetraploid cells (Fig. 7c), suggesting that cytokinesis *per se*, in which BRCA2 is also involved ^11–13^, is not affected.

Finally, the function of BRCA2-PLK1 interaction in mitosis seems to be independent of the HR function of BRCA2 as cells expressing these variants display mild sensitivity to DNA damage (MMC) and PARP inhibitors (Fig. 8a-c), normal recruitment of RAD51 to DNA damage sites (Fig. 8e, Supplementary Fig. 8b), absence of micronuclei (Supplementary Fig. 8d, e), rescue the chromosome bridges phenotype of BRCA2 deficient cells (Figure 6d) and their HR activity, as measured by DSB-mediated gene targeting, is intact (Fig. 8f, Supplementary Fig. 9). Nevertheless, we cannot rule out that the mild phenotype observed in some of these experiments in cells bearing S206C and T207A may arise from a distorted interaction with an unknown DNA repair factor that would bind to the region of BRCA2 where these variants localize.

Putting our results together we reveal an unexpected chromosome stability control mechanism that depends on the phosphorylation of BRCA2 by PLK1 at T207. We show that BRCA2 pT207 is a docking platform for PLK1 that ensures the efficient interaction of BUBR1 with PP2A phosphatase required for chromosome alignment.

We propose the following working model (Fig. 8g): in cells expressing BRCA2 WT, PLK1 phosphorylates BRCA2 on T207 leading to the docking of PLK1 at this site. This step promotes the formation of a complex between BRCA2-PLK1-pBUBR1-PP2A in prometaphase at the kinetochore, leading to an enrichment of phosphorylated BUBR1 and the phosphatase PP2A to balance Aurora B activity (Fig. 8g, panel 1). Why is BRCA2 required for PP2A interaction with pBUBR1 and, is this regulated by PLK1? It has been reported that BRCA2 fragment from aa 1001 to aa 1255 (BRCA2_1001-1255_) comprising a PP2A-B56 binding motif, binds to the B56 subunit of PP2A^41^; in addition, the phosphorylation of positions 2 and 8 of this motif enhances the binding to B56^42^. In BRCA2, these positions are occupied by serines that are targets for PLK1. Therefore, it is likely that the recruitment of PLK1 by BRCA2 T207 would favor phosphorylation of BRCA2_1001-1255_ and binding of the complex between BRCA2 and PLK1 to PP2A-B56. Non-exclusively, BUBR1 has been reported to bind directly BRCA2 although there are inconsistencies regarding the site of interaction ^4, 5^. Thus, either the direct interaction of BRCA2 with PP2A or with BUBR1 or both could be favoring the formation of a complex between PP2A and pBUBR1.

Once the kinetochore-microtubule attachments are established and proper tension is achieved, BUBR1 is no longer phosphorylated by PLK1^19^. This leads to a full alignment of chromosomes at the metaphase plate, SAC inactivation (Fig. 8g, panel 2) and the subsequent faithful chromosome segregation in anaphase (Fig. 8g, panel 3).

In cells expressing the variants that impair T207 phosphorylation (S206C, T207A), PLK1 cannot be recruited to pT207-BRCA2, impairing the formation of the complex with PLK1, PP2A and BUBR1 (Fig. 8g, panel 1’), which in turn reduces the amount of pBUBR1 and its binding to PP2A for proper kinetochore-microtubule interactions. This leads to chromosome misalignment defects that prolong mitosis (Fig. 8g, panel 2’); as a consequence, these cells exhibit increased chromosome segregation errors (Fig. 8g, panel 3’) and aneuploidy.

Although the individual BRCA2 variants analysed here are rare (Supplementary table 1), the majority of pathogenic mutations recorded to date lead to a truncated protein either not expressed or mislocalized ^43^ which would be predicted to affect this function. Consistent with this idea, the BRCA2 deficient cells or cells transiently depleted of BRCA2 used in this study exhibit low levels of phosphorylated BUBR1 (Fig. 5f, g). Thus, the chromosome alignment function described here could be responsible, at least in part, for the numerical chromosomal aberrations observed in *BRCA2*-associated tumors ^44^.

Finally, the lack of sensitivity to the PARP inhibitor Olaparib observed in our cell lines (Fig. 8b) has important clinical implications as breast cancer patients carrying these variants are not predicted to respond to PARP inhibitor treatment (unlike *BRCA2*-mutated tumors that are HR-deficient).

## Supporting information

3D structure PLK1_PBD-BRCA2pT207

Report 3D structure

## Acknowledgments

We thank members of the AC lab for fruitful comments on the manuscript and Davide Panigada and Jose M. Jimenez-Gomez for the illustration of chromosomes in Figure 8. We thank Rene H. Medema for useful discussions on this work including the cell synchronization protocol used in Figure 7. We also thank Juan S. Martinez for construct BRCA21-250, Anne Houdusse for construct PLK1365-603, Carine Giovannangeli for TALEN plasmids, Eric Nigg for pS676-BUBR1 antibody and Geert JPL Kops for BUBR1-RFP construct. We acknowledge the Cell and Tissue Imaging Facility of the Institut Curie (PICT), a member of the France BioImaging National Infrastructure (ANR-10-INBS-04), and the French Infrastructure for Integrated Structural Biology (https://www.structuralbiology.eu/networks/frisbi, ANR-10-INSB-05-01). We thank Charlene Lasgi from the Flow Cytometry platform of Institut Curie, Orsay. We thank Guillaume Hoffmann and Jose A. Marquez from the HTXLab (Grenoble, France), funded by the European Community’s Seventh Framework Programme H2020 under iNEXT (grant agreement N°653706).

This work was supported by the ATIP-AVENIR CNRS/INSERM Young Investigator grant 201201, FRM ‘Amorcage Jeunes Equipes’ Young Investigator grant AJE20110, EC-Marie Curie Career Integration grant CIG293444 to A.C. and Institut National du Cancer INCa-DGOS_8706 to A.C. and S.Z.J.; A.E. was supported by the Swedish Society for Medical Research.

## Author Contributions

A.E. purified WT and mutated BRCA2_1-250_, established the stable DLD1^-/-^ cell lines, performed kinase assays, pull-down assays, Western blots, time-lapse microscopy experiments, mitotic index measurements by FACS, clonogenic survival and MTT assays as well as the statistical analysis for all the experiments. C.M. performed IF and image acquisition of metaphase plate alignment, chromosome segregation and karyotype analysis. M.J. performed the NMR experiments assisted by S. M., F.T. and S.Z.J. M.J. and S.M. purified PLK1_PBD_ and performed the ITC experiments. S.M. solved the X-ray structure assisted by V.R. R.B. and G.S. performed IF, image acquisition and quantification of DNA repair foci. V.B. assisted establishing stable clones and performing clonogenic survival assays. P.D. purified PLK1_PBD_. A.M. cloned and produced PLK1 and PLK1-KD from insect cells. A.C., A.E. and S.Z.J. designed the experiments. A.C. and S.Z.J. supervised the work. A.C. wrote the paper with important contributions from all authors.

The authors declare no conflict of interest.

## References

1. Moynahan, M. E., Pierce, A. J. & Jasin, M. BRCA2 is required for homology-directed repair of chromosomal breaks. Molecular Cell 7, 263–272 (2001).

2. Jensen, R. B., Carreira, A. & Kowalczykowski, S. C. Purified human BRCA2 stimulates RAD51-mediated recombination. Nature 467, 678–683 (2010).

3. Saleh-Gohari, N. & Helleday, T. Conservative homologous recombination preferentially repairs DNA double-strand breaks in the S phase of the cell cycle in human cells. Nucleic Acids Research 32, 3683–3688 (2004).

4. Choi, E. et al. BRCA2 fine-tunes the spindle assembly checkpoint through reinforcement of BubR1 acetylation. Developmental Cell 22, 295–308 (2012).

5. Futamura, M. et al. Potential role of BRCA2 in a mitotic checkpoint after phosphorylation by hBUBR1. Cancer Res 60, 1531–1535 (2000).

6. Lampson, M. A. & Kapoor, T. M. The human mitotic checkpoint protein BubR1 regulates chromosome-spindle attachments. Nat Cell Biol 7, 93–98 (2005).

7. Lara-Gonzalez, P., Westhorpe, F. G. & Taylor, S. S. The Spindle Assembly Checkpoint. Current Biology 22, R966–R980 (2012).

8. Elowe, S. et al. Uncoupling of the spindle-checkpoint and chromosome-congression functions of BubR1. J Cell Sci 123, 84–94 (2010).

9. Zhang, G., Mendez, B. L., Sedgwick, G. G. & Nilsson, J. Two functionally distinct kinetochore pools of BubR1 ensure accurate chromosome segregation. Nature Communications 7, 12256 (2016).

10. Park, I. et al. HDAC2/3 binding and deacetylation of BubR1 initiates spindle assembly checkpoint silencing. FEBS J 284, 4035–4050 (2017).

11. Mondal, G. et al. BRCA2 localization to the midbody by filamin A regulates cep55 signaling and completion of cytokinesis. Developmental Cell 23, 137–152 (2012).

12. Daniels, M. J., Wang, Y., Lee, M. & Venkitaraman, A. R. Abnormal cytokinesis in cells deficient in the breast cancer susceptibility protein BRCA2. Science 306, 876–879 (2004).

13. Takaoka, M., Saito, H., Takenaka, K., Miki, Y. & Nakanishi, A. BRCA2 phosphorylated by PLK1 moves to the midbody to regulate cytokinesis mediated by nonmuscle myosin IIC. Cancer Res 74, 1518–1528 (2014).

14. Lin, H.-R., Ting, N. S. Y., Qin, J. & Lee, W.-H. M phase-specific phosphorylation of BRCA2 by Polo-like kinase 1 correlates with the dissociation of the BRCA2-P/CAF complex. J Biol Chem 278, 35979–35987 (2003).

15. Lee, M., Daniels, M. J. & Venkitaraman, A. R. Phosphorylation of BRCA2 by the Polo-like kinase Plk1 is regulated by DNA damage and mitotic progression. Oncogene 23, 865–872 (2004).

16. Zitouni, S., Nabais, C., Jana, S. C., Guerrero, A. & Bettencourt-Dias, M. Polo-like kinases: structural variations lead to multiple functions. Nat Rev Mol Cell Biol 15, 433– 452 (2014).

17. Barr, F. A., Silljé, H. H. W. & Nigg, E. A. Polo-like kinases and the orchestration of cell division. Nat Rev Mol Cell Biol 5, 429–440 (2004).

18. Huang, H. et al. Phosphorylation sites in BubR1 that regulate kinetochore attachment, tension, and mitotic exit. The Journal of Cell Biology 183, 667–680 (2008).

19. Elowe, S., Hümmer, S., Uldschmid, A., Li, X. & Nigg, E. A. Tension-sensitive Plk1 phosphorylation on BubR1 regulates the stability of kinetochore microtubule interactions. Genes Dev 21, 2205–2219 (2007).

20. Suijkerbuijk, S. J. E., Vleugel, M., Teixeira, A. & Kops, G. J. P. L. Integration of kinase and phosphatase activities by BUBR1 ensures formation of stable kinetochore-microtubule attachments. Developmental Cell 23, 745–755 (2012).

21. Hauf, S. et al. The small molecule Hesperadin reveals a role for Aurora B in correcting kinetochore-microtubule attachment and in maintaining the spindle assembly checkpoint. J Cell Biol 161, 281–294 (2003).

22. Foley, E. A., Maldonado, M. & Kapoor, T. M. Formation of stable attachments between kinetochores and microtubules depends on the B56-PP2A phosphatase. Nat Cell Biol 13, 1265–1271 (2011).

23. Elia, A. E. H., Cantley, L. C. & Yaffe, M. B. Proteomic screen finds pSer/pThr-binding domain localizing Plk1 to mitotic substrates. Science 299, 1228–1231 (2003).

24. Elia, A. E. H. et al. The molecular basis for phosphodependent substrate targeting and regulation of Plks by the Polo-box domain. Cell 115, 83–95 (2003).

25. Neef, R. et al. Phosphorylation of mitotic kinesin-like protein 2 by polo-like kinase 1 is required for cytokinesis. J Cell Biol 162, 863–875 (2003).

26. Kang, Y. H. et al. Self-regulated Plk1 recruitment to kinetochores by the Plk1-PBIP1 interaction is critical for proper chromosome segregation. Mol. Cell 24, 409–422 (2006).

27. Szabo, C., Masiello, A., Ryan, J. F. & Brody, L. C. The breast cancer information core: database design, structure, and scope. Hum. Mutat. 16, 123–131 (2000).

28. Béroud, C. et al. BRCA Share: A Collection of Clinical BRCA Gene Variants. Hum. Mutat. 37, 1318–1328 (2016).

29. Golsteyn, R. M., Mundt, K. E., Fry, A. M. & Nigg, E. A. Cell cycle regulation of the activity and subcellular localization of Plk1, a human protein kinase implicated in mitotic spindle function. J Cell Biol 129, 1617–1628 (1995).

30. Yun, S.-M. et al. Structural and functional analyses of minimal phosphopeptides targeting the polo-box domain of polo-like kinase 1. Nat Struct Mol Biol 16, 876–882 (2009).

31. García-Alvarez, B., de Cárcer, G., Ibañez, S., Bragado-Nilsson, E. & Montoya, G. Molecular and structural basis of polo-like kinase 1 substrate recognition: Implications in centrosomal localization. Proc Natl Acad Sci USA 104, 3107–3112 (2007).

32. Hucl, T. et al. A syngeneic variance library for functional annotation of human variation: application to BRCA2. Cancer Res 68, 5023–5030 (2008).

33. Yata, K. et al. BRCA2 coordinates the activities of cell-cycle kinases to promote genome stability. CellReports 7, 1547–1559 (2014).

34. Lera, R. F. et al. Decoding Polo-like kinase 1 signaling along the kinetochore-centromere axis. Nature Chemical Biology 12, 411–418 (2016).

35. Wang, J. et al. Crystal structure of a PP2A B56-BubR1 complex and its implications for PP2A substrate recruitment and localization. Protein Cell 7, 516–526 (2016).

36. Santaguida, S. & Amon, A. Short-and long-term effects of chromosome mis-segregation and aneuploidy. Nat Rev Mol Cell Biol 16, 473–485 (2015).

37. Kraakman-van der Zwet, M. et al. Brca2 (XRCC11) deficiency results in radioresistant DNA synthesis and a higher frequency of spontaneous deletions. Mol. Cell. Biol. 22, 669–679 (2002).

38. Bryant, H. E. et al. Specific killing of BRCA2-deficient tumours with inhibitors of poly(ADP-ribose) polymerase. Nature 434, 913–917 (2005).

39. Brunet, E. et al. Chromosomal translocations induced at specified loci in human stem cells. Proc Natl Acad Sci USA 106, 10620–10625 (2009).

40. Hockemeyer, D. et al. Efficient targeting of expressed and silent genes in human ESCs and iPSCs using zinc-finger nucleases. Nat Biotechnol 27, 851–857 (2009).

41. Hertz, E. P. T. et al. A Conserved Motif Provides Binding Specificity to the PP2A-B56 Phosphatase. Mol. Cell 63, 686–695 (2016).

42. Wu, C.-G. et al. PP2A-B’ holoenzyme substrate recognition, regulation and role in cytokinesis. Cell Discov 3, 17027–19 (2017).

43. Spain, B. H., Larson, C. J., Shihabuddin, L. S., Gage, F. H. & Verma, I. M. Truncated BRCA2 is cytoplasmic: implications for cancer-linked mutations. Proc Natl Acad Sci USA 96, 13920–13925 (1999).

44. Gretarsdottir, S. et al. BRCA2 and p53 Mutations in Primary Breast Cancer in Relation to Genetic Instability. Cancer Res 58, 859–862 (1998).

