## Supplementary tables, suppl. legends 1-10 for "Proper chromosome alignment depends on BRCA2 phosphorylation by PLK1"

**Supplementary information**

**Supplementary tables 1-2, Supplementary Figures and Supplementary Figure legends 1-10**

**Supplementary table 1:** Number of records in ClinVar and BRCAShare data bases of the variants identified in breast cancer patients altering the amino acids investigated in this work. The specific VUS utilized in this study are highlighted in bold.

| **VUS** | **ClinVar** | **BRCAShare** |
| --- | --- | --- |
| **M192T** | 2 | 0 |
| S196I | 4 | 0 |
| **S196N** | 4 | 3 |
| S196T | 1 | 0 |
| **S206C** | 1 | 1 |
| S206Y | 1 | 0 |
| **T207A** | 3 | 0 |
| T207I | 4 | 0 |

**Supplementary table 2:** Statistics table for the crystal structure of PBD_pT207, a complex between the Polo-Box Domain of human PLK1 (aa 365 to aa 603) and the 17aa peptide pT207 of BRCA2 (aa 194 to aa 210, threonine 207 being phosphorylated).

|  | PBD_T207 phosphorylated  (PDB 6GY2) |
| --- | --- |
| **Data collection** |  |
| Space group | P_1_ |
| Cell dimensions : |  |
| *a*, *b*, *c* (Å) | 50.000 56.040 61.030 |
| α, β, γ (°) | 80.79 79.23 65.05 |
| Molecules per a.u  Resolution (Å) | 2  59.70 – 3.106 (3.106 – 3.16) |
| *R*_merge_ | 0.056 (0.346) |
| *R*_meas_ | 0.076 (0.466) |
| *R*_pim_ | 0.051 (0.31) |
| *I/*σ (*I*) | 10.1 (2.3) |
| *CC*_1/2_ | 0.997 (0.821) |
| Completeness (spherical, %) | 93.7 (96.9) |
| Redundancy | 1.94 (1.977) |
| B Wilson (Å²)  Multiplicity | 76.9 (78.86)  1.9 (2.0) |
| **Refinement** |  |
| Resolution (Å) | 22.57 - 3.11 (3.11 – 3.15) |
| No. reflections | 9911 |
| *R*_work_ / *R*_free_ | 0.189/0.215 |
| No. Atoms |  |
| Protein | 3788 |
| Heterogen atoms | 28 |
| Water | 22 |
| R.m.s. deviations |  |
| Bond lengths (Å) | 0.007 |
| Bond angles (°) | 0.97 |

*Values in parentheses are for highest-resolution shell.

**Supplementary Figure 1**. Related to Figure 1. **PLK1 phosphorylation of the N-terminal region of BRCA2 and conservation of PLK1 phosphosites**

**(a)** PLK1 *in vitro* kinase assay with BRCA2_1-250_. Top: The polypeptide 2x-MBP-BRCA2_1-250_ WT was incubated with increased concentrations (0, 50 and 100 ng) of recombinant PLK1 in the presence of γ^32^P-ATP. The samples were resolved on 4-15% SDS-PAGE and the ^32^P-labeled products were detected by autoradiography. Bottom: 4-15% SDS-PAGE showing the input of 2xMBP-BRCA2_1-250_ WT (0.5 μg) used in the reaction. **(b)** Quantification of the relative phosphorylation in (a). Data are represented as mean ± SD from two independent experiments. One-way ANOVA test with Dunnett’s multiple comparisons test was used to calculate statistical significance of differences. **(c)** Alignment of the region 2-284 of human BRCA2 with the N-terminal regions of BRCA2 from 40 different species. Amino acids conserved in more than 30 % of the species are highlighted with coloured background. A dashed line box identifies the highly conserved cluster around S193 (amino acid 180 to amino acid 210). Arrows show the amino acids cited in this manuscript (including the PLK1 phosphosites).

**Supplementary Figure 2.** Related to Figure 2. **PLK1 phosphorylation kinetics of BRCA2_190-284_ WT vs T207A**

**(a)** Superimposition of the ^1^H-^15^N HSQC spectra recorded on the ^15^N labelled fragments BRCA2_190-284_ WT (red) and T207A (green) after 5h of incubation with PLK1. The conditions are the same as in Figure 1. **(b,c)** Comparison between the phosphorylation kinetics of BRCA2_190-284_ WT and T207A is displayed for 2 different experiments performed with 2 different PLK1 samples. The percentage of phosphorylation deduced from the intensities of the peaks corresponding to the non-phosphorylated and phosphorylated residues is plotted as a function of time. WT S193 and T207 time points are represented by red circles and squares, respectively, while T207A S193 timepoints are represented by green circles.

**Supplementary Figure 3.** Related to Figure 3. **Isothermal Titration Calorimetry (ITC) thermogram showing binding of PLK1_PBD_ to the fragment BRCA2_190-284_ or a 10 aa BRCA2 peptide containing pS197**

Thermogram showing the binding affinity of PLK1_PBD_ to the **(a)** phosphorylated or **(b)** non-phosphorylated BRCA2_190-284_ fragment, purified from bacteria as explained in the Methods section and also used in the NMR experiments. **(c)** Thermogram showing the binding affinity of PLK1_PBD_ to a 17 aa BRCA2 synthetic peptide comprising T207D (WSSSLATPPTLSSD_207_VLI). **(d)** Thermogram showing the binding affinity of PLK1_PBD_ to a 10 aa BRCA2 synthetic peptide comprising pS197.

**Supplementary Figure 4.** Related to Figures 3-8**. BRCA2 protein levels in DLD1 BRCA2^-/-^ stable clones complemented with the cDNA of BRCA2 WT and variants utilized in this study and effect of PLK1 and CDK1 inhibitors on the interaction between BRCA2 and PLK1**

**(a)** BRCA2 protein levels in total protein extracts from the DLD1 BRCA2 deficient (BRCA2^-/-^) cells stably expressing GFP-MBP-BRCA2 WT (BRCA2 WT C1) or the variants S206C (clones A7 and A9) and T207A (clones B1 and E4) as detected by western blot using anti-BRCA2 (OP95) antibody. **(b)** The effect of PLK1 inhibitors on the interaction between GFP-MBP-BRCA2 (BRCA2 WT) and PLK1. BRCA2 WT cells were treated with nocodazole (100 ng/ml) for 14h before the PLK1 inhibitor BTO (50 µM) was added to the cells followed by additional two hours incubation before being harvested. The cells were lysed and GFPMBP-BRCA2 was immunoprecipitated with GFP-trap beads, immunocomplexes were resolved on 4-15% SDS-PAGE followed by western blotting, the interaction with PLK1 was revealed by anti-PLK1 and -MBP antibodies. Unsynchronized DLD1 cells with endogenous BRCA2 (BRCA2^+/+^) were used as control for the immunoprecipitation and StainFree images of the gels before transfer were used as loading control for the input (cropped image is shown). The amount of PLK1 co-immunoprecipitated with GFPMBP-BRCA2 relative to the input levels of PLK1 and the amount of immunoprecipitated GFPMBP-BRCA2 is presented below the blot, relative to non-treated BRCA2 WT. **(c)** The effect of CDK1 inhibitors on the interaction between 2xMBP-BRCA21-250 and PLK1. U2OS cells were transient transfected with the 2xMBP-BRCA21-250 WT construct, nocodazole (300 ng/ml) was added to the cells 30h post-transfection and the cells were treated for 14h before CDK1 inhibitor (Ro-3306 50nM) was added followed by two hours additional incubation. The cells were lysed and immunoprecipitation was performed against the MBP tag using amylose beads. Complexes were resolved on 4-15% SDS-PAGE followed by western blotting using anti-PLK1 and anti-MBP antibodies. The amount of PLK1 co-immunoprecipitated with 2xMBP-BRCA2_1-250_ relative to the input levels of PLK1 and the amount of immunoprecipitated 2xMBP-BRCA2_1-250_ is presented below the blot as mean ± SD from three independent experiments. The data is presented relative to the non-treated BRCA2 WT.

**Supplementary Figure 5.** Related to Figure 5**. BRCA2 at the kinetochores and effect of PLK1 inhibitors and phosphatase treatment on pT680-BUBR1.**

**(a)** Representative images of the localization of BRCA2 in nocodazole-arrested U2OS transient expressing GFP-MBP-BRCA2 WT. BRCA2 is detected by anti-BRCA2 rabbit antibody (CA1033), CREST is used as centromere marker and DNA is counterstained with DAPI. Scale bar represents 1 µm. **(b)** Protein levels of pT680-BUBR1 in BRCA2 WT stable clone after treatment with PLK1 inhibitors. After 14h culture with media containing nocodazole (100 ng/µl), PLK1 inhibitors (Bi2536 (50 nM) or BTO (50 µM)) were added to the media and the cells were cultured for additional 2h before harvesting. The level of pT680-BUBR1 was analyzed in the total protein extract by western blot. **(c)** Phosphatase (Fast AP phosphatase) treatment of total protein lysate extracted from DLD1 BRCA2 WT cells treated with nocodazole (100 ng/µl) for 14h. **(d)** Western blot showing the expression levels of endogenous BUBR1 and pS676-BUBR1 in nocodazole treated stable clones of DLD1 BRCA2 deficient cells (BRCA2^-/-^) expressing GFPMBP-BRCA2 WT (BRCA2 WT) or T207A variant. The levels of pS676-BUBR1 was analyzed in the total protein extract by western blot. The mean pBUBR1 signal relative to the stain free signal is shown for the nocadozole treated samples below the blots. Results are presented normalized to the protein levels for BRCA2 WT. The data represents the mean ± SD of two independent experiments.

**Supplementary Figure 6**. Related to Figure 5. **Localization of BRCA2 and PLK1 at the kinetochores and PLK1-BUBR1 complex in cells bearing variant T207A compared to WT**

**(a)** Representative images of the localization of PLK1 in nocodazole-arrested DLD1 BRCA2^-/-^ cells stably expressing GFP-MBP-BRCA2 WT or the variant T207A as indicated. CREST is used as centromere marker and DNA is counterstained with DAPI. Scale bar represents 1 µm. **(b)** Quantification of the co-localization of PLK1 and CREST in (a). The data represents the intensity ratio (PLK1:CREST) relative to the mean ratio of PLK1:CREST for the GFP-MBP-BRCA2 WT calculated from a total of 180 pairs of chromosomes analysed from two independent experiments (6 pairs of chromosomes/cell from 15 cells). The red line in the plot indicates the median (95% CI) ratio, each dot represents a pair of chromosomes. For statistical comparison of the differences between the samples we applied a Mann-Whitney test, the p-values show significant difference. **(b)** Co-immunoprecipitation of endogenous PLK1 with endogenous BUBR1 from mitotic cell extracts of BRCA2 WT cells or cells expressing the variants S206C or T207A using mouse anti-BUBR1 antibody. Mouse IgG was used as control for the BUBR1 immunoprecipitation. The immuno-complexes were resolved on 4-15% SDS-PAGE followed by western blotting, the interactions were revealed by rabbit anti-BUBR1 and mouse anti-PLK1 antibodies. **(d)** Quantification of co-immunoprecipitated PLK1 in (c), relative to the input levels and the amount of immunoprecipitated BUBR1. Results are presented as the fold change compared to the BRCA2 WT clone. The data represents the mean ± SD of three to four independent experiments. Statistical significance of the difference was calculated with one-way ANOVA test with Dunnett’s multiple comparisons test, the p-values show the significant difference, ns: non-significant.

**Supplementary Figure 7.** Related to Figure 7**. BrdU incorporation measured by flow cytometry of DLD1 BRCA2^-/-^ stable cell lines expressing BRCA2 WT or BRCA2 variants S206C and T207A.**

**(a-c)** Representative flow cytometry plots for the analysis of S-phase tetraploid cells (quantified in Figure 7c, d) in the stable DLD1 BRCA2 deficient cells expressing BRCA2 WT (a) or the VUS S206C (b) and T207A (c). Viable cells were gated from the Forward Scatter (FSC-A) versus Side Scatter (SSC-A) plots and displayed in a 7-AAD-W versus 7-AAD-A plot to exclude doublets. The gated singlet population was displayed in a APC-A (BrdU) versus 7-AAD-A (DNA) plot. The S-phase tetraploid population was gated as BrdU^+^ cells with DNA content >4N. 20 000 singlet events were collected for each experiment. **(d)** Frequency of BrdU^+^ cells in the stable clones expressing BRCA2 WT or the VUS as indicated. The data represents the mean ± SD of three independent experiments. Statistical significance of the difference was calculated with one-way ANOVA test with Tukey’s multiple comparisons test (the p value show the difference compared to WT, ns: non-significant).

**Supplementary Figure 8.** Related to Figure 8**. Plating efficiency of unchallenged DLD1 BRCA2^-/-^ stable clones expressing EGFP-MBP-BRCA2 WT or the variants. and representative images of DNA damage foci in these cells. Micronuclei in BRCA2^-/-^ stable cell lines and BRCA2^-/-^ expressing BRCA2 WT or BRCA2 variants S206C and T207A.**

**(a)** Representative plates showing the number of colonies in unchallenged conditions of the cells assessed for MMC-clonogenic survival assay quantified in Fig. 8a (500 cells seeded per 6-well plate). **(b)** Representative immunofluorescence images of nuclear γH2AX and RAD51 foci in DLD1 BRCA2^+/+^ cells depleted of BRCA2 (siBRCA2), BRCA2^+/+^control cells (siCTRL), DLD1 BRCA2 deficient cells (BRCA2^-/-^) stably expressing BRCA2 WT or the variant T207A, in non-treated (-IR) or two hours after exposure to 6 Gy of γ-irradiation (+IR), as indicated in the images and as quantified in Fig. 8d, 8e. Scale bar represents 10 µm. **(c)** Western Blot showing the levels of endogenous BRCA2 in the siRNA transfected cells imaged in (b) and analysed in Fig. 8d, 8e, at the time for radiation. **(d)** Frequency of micronuclei with and without centromeres in DLD1 BRCA2 deficient cells (BRCA2^-/-^) and BRCA2^-/-^ clones stably expressing BRCA2 WT or the variants S206C and T207A. *n* indicates the total number of cells counted for each clone from two independent experiments. Statistical significance of the difference was calculated with two-way ANOVA test with Tukey’s multiple comparisons test, the p-values show the significant difference, ns: non-significant. **(e)** Representative images of micronuclei with and without centromeres observed in cells quantified in (d), scale bar represents 10 µm.

**Supplementary Figure 9.** Related to Figure 8f**. Frequency of mCherry positive cells measured by flow cytometry of DLD1 BRCA2^-/-^ stable cell lines expressing BRCA2 WT or BRCA2 variants S206C and T207A.**

**(a-e)** Representative flow cytometry plots for the analysis of mCherry positive cells in the HR assay (quantified in Fig. 8f) in DLD1 cells with endogenous BRCA2 (BRCA2^+/+^) (a), DLD1 BRCA2 deficient cells (BRCA2^-/-^) (b), BRCA2 deficient cells expressing BRCA2 WT (c) or the VUS S206C (d) and T207A (e). Viable cells were gated from the Forward Scatter (FSC-A) versus Side Scatter (SSC-A) plots and displayed in a SSC-H versus SSC-A plot to exclude doublets. The gated singlet population was displayed in a mCherry-A versus FSC-A plot. The mCherry positive population was gated from non-transfected cells. 10 000 singlet events were collected for each experiment.

**Supplementary Figure 10. SDS-PAGE of the PLK1, PLK1-KD and PLK1-PBD recombinant proteins utilized in this study and comparison of the kinase activity of each batch of PLK1**

**(a)** SDS-PAGE showing purified PLK1, PLK1-K82R mutant (PLK1-KD) and PLK1_PBD_. Human PLK1 was expressed and purified from sf9 insect cells using Ni-NTA column followed by a second purification step with a cationic exchange Capto S column. Purified PLK1 and PLK1-K82R protein (3 µg) were loaded on a 4-15% SDS-PAGE Stain-Free gel. For purification of PLK1_PBD,_ 6His-Sumo-PLK1_PBD_ was expressed and purified from bacteria using a His-TRAP column, the His-tag was cleaved with 6xHis-SUMO Protease and the cleaved PLK1_PBD_ was further purified using Ni-NTA agarose resin. The purified protein was loaded on a 4-20% SDS-PAGE (1.4 µg) and detected by Coomassie staining. **(b)** *In vitro* kinase assay with the purified PLK1 (0.1 µg) from (a) or PLK1 purchased from Abcam, 0.1 µg PLK1 was used in the kinase reaction with either RAD51 (25 ng) or purified 2xMBP-BRCA2_1-250_ WT (0.5 µg) as substrate in the presence of [γ^32^P]-ATP. The samples were resolved by 7.5 % SDS-PAGE and ^32^P-labeled products were detected by autoradiography.
