## Supplementary material for "Proper chromosome alignment depends on BRCA2 phosphorylation by PLK1": Materials and Methods

**Cell lines, cell culture and synchronisations**

The human cell lines HEK293T and U2OS cells (kind gift from Dr. Mounira Amor-Gueret) were cultured in DMEM (Eurobio Abcys, Courtaboeuf, France) media containing 25 mM sodium bicarbonate and 2 mM L-Glutamine supplemented with 10% heat inactive FCS (EuroBio Abcys). The BRCA2 deficient colorectal adenocarcinoma cell line DLD1 BRCA2^-/-^ (Hucl, T. et al 2008) (HD 105-007) and the parental cell line DLD1 BRCA2^+/+^ (HD-PAR-008) was purchased from Horizon Discovery (Cambridge, England). The cells were cultured in RPMI media containing 25 mM sodium bicarbonate and 2 mM L-Glutamine (EuroBio Abcys) supplemented with 10% heat inactive FCS (EuroBio Abcys). The DLD1 BRCA2^-/-^ cells were maintained in growth media containing 0.1 mg/ml hygromycin B (Thermo Fisher Scientific). The stable cell lines of DLD1^-/-^ BRCA2 deficient cells expressing BRCA2 WT or variants of interest generated in this study were cultured in growth media containing 0.1 mg/ml hygromycin B and 1 mg/ml G418 (Sigma-Aldrich). All cells were cultured at 37°C with 5% CO_2_ in a humidified incubator and all cell lines used in this study have been regularly tested negatively for mycoplasma contamination.

For synchronization of cells in mitosis, nocodazole (100-300 ng/ml, Sigma-Aldrich) was added to the growth media and the cells were cultured for 14h before harvesting. For synchronisation by double thymidine block, the cells were treated with thymidine (2.5 mM, Sigma-Aldrich) for 17h, released for 8h followed by a second thymidine (2.5 mM) treatment for 15h.

**Plasmids**

2XMBP-, human 2XMBP-BRCA2_1-250_ and EGFP-MBP-BRCA2 subcloning in phCMV1 expression vector were generated as described ^49,50^. In the case of 2XMBP and 2XMBP-BRCA2_1-250_, a tandem of 2 nuclear localization signals from RAD51 sequence was added downstream the MBP-tag.

Point mutations (M192T, S193A, S196N, S206C and T207A) were introduced in the 2xMBP-BRCA2_1-250_, EGFP-MBP-BRCA2 vector using QuikChange II and QuikChange XL site-directed mutagenesis kit (Agilent Technologies), respectively (see Table 1, 2 for primer sequences).

For expression of BRCA2_48-218_ in bacteria, an optimized gene coding for human His-tagged BRCA2_48-218_ (WT and T207A) was synthetized by Genscript and cloned in a pETM13 vector (a TEV site being present between the tag and the BRCA2 fragment). For expression of BRCA2_190-284_ in bacteria, the human BRCA2_190-284_ was amplified by PCR using full length BRCA2 as template (phCMV1-2xMBP-BRCA2, see Table 3 for primer sequences). The PCR product was purified and digested with *BamH1* and *SalI* and cloned into in the pGEX-6P-1 vector (GE Healthcare) to generate GST-BRCA2_190-284_. The point mutation T207A was introduced in the same way in BRCA2_190-284_ as in 2xMBP-BRCA2_1-250_ and the EGFP-MBP-BRCA2. The introduction of the point mutation was verified by sequencing (see Table 1 and Table 2 for primer sequences).

The *PLK1* cDNA (Addgene pTK24) was cloned into the pFast-Bac HT vector using Gibson assembly (NEB) (see Table 4 for primer sequences). To produce PLK1-KD, the point mutation K82R was introduced in the pFast-Bac HT-PLK1 vector using QuikChange XL site-directed mutagenesis kit (see Table 5 for primer sequences).

The Polo-like binding domain (PBD) of PLK1 (amino acid 326 to amino acid 603) was amplified from the pTK24 plasmid (Addgene) and cloned into a pT7-His6-SUMO expression vector using NEB Gibson assembly (Gibson Assembly Master Mix, New England BioLabs, Cat. # E2611S) (see Table 5 for primer sequences). A plasmid containing a smaller PLK1 PBD fragment (amino acid 365 to amino acid 603) with a N-terminal GST tag was a kind gift from Dr. Anne Houdusse (Institute Curie, Paris).

To produce the phosphomimic BUBR1 mutant, we introduced the S670D, S676D and T680D point mutations in the pcDNA3-3xFLAG-BUBR1-RFP construct (kind gift from Dr. Geert JPL Kops) using QuikChange XL site-directed mutagenesis kit (see Table 6 for primer sequences).

For the DSB-gene targeting assay, we replaced the GFP tag in the promoter-less AAVS1-2A-GFP-pA plasmid (kind gift from Dr. Carine Giovannangeli) with the mCherry tag from the pET28 mCherry plasmid using NEB Gibson Assembly (Gibson Assembly Master Mix, New England BioLabs, Cat. # E2611S). See Table 9 for primer sequences.

**Expression and purification of 2xMBP-BRCA2_1-250_**

The 2xMBP-BRCA2_1-250_ was purified as previously described^49^. Briefly, ten 150 mm plates of HEK293T were transient transfected with the 2xMBP-BRCA2_1-250_ using TurboFect (Thermo Fisher Scientific). The cells were harvested 30 h post transfection, lysed in lysis buffer H (50 mM HEPES (pH 7.5), 250 mM NaCl, 1% NP-40, 5 mM EDTA, 1 mM DTT, 1 mM PMSF and EDTA-free Protease Inhibitor Cocktail (Roche)) and incubated with amylose resin (NEB) for 3h at 4°C. The 2xMBP-BRCA2_1-250_ was eluted with 10 mM maltose. The eluate was further purified with Bio-Rex 70 cation-exchange resin (Bio-Rad) by NaCl step elution. The size and purity of the final fractions were analysed by SDS-PAGE and western blotting using anti-MBP antibody. The 2xMBP-BRCA2_1-250_ fragments containing the BRCA2 variants (M192T, S193A, S196N, S206C and T207A) were purified following the same protocol as for WT 2xMBP-BRCA2_1-250_.

**Expression and purification of BRCA2_48-218_ and BRCA2_190-284_ for NMR**

Recombinant ^15^N-labelled (WT, T207A) and ^15^N/^13^C-labelled (WT) BRCA2_48-218_were produced by transforming *Escherichia coli* BL21 (DE3) Star cells with the pETM13 vector containing human BRCA2_48-218_ (WT and T207A). Recombinant ^15^N-labelled (WT, T207A) and ^15^N/^13^C-labelled (WT, T207A) BRCA2_190-284_ were produced by transforming *Escherichia coli* BL21 (DE3) Star cells with the pGEX-6P-1 vector containing human BRCA2_190-284_ (WT and T207). Cells were grown in a M9 medium containing 0.5 g/l ^15^NH_4_Cl and 2 g/l ^13^C-glucose when ^13^C labelling was needed. The bacterial culture was induced with 1 mM IPTG at an OD_600_ of 0.8, and it was further incubated for 3 h at 37°C. Harvested cells were resuspended in buffer A (50 mM Tris-HCl pH 8.0, 150 mM NaCl, 2 mM DTT, 1 mM EDTA) with 1 mM PMSF and 1X protease inhibitors cocktail (Roche) and disrupted by sonication. For BRCA2_48-218_, clarified cell lysate was loaded onto Ni-NTA beads (Thermo Scientific) equilibrated with buffer A. After 1 h of incubation at 4°C, beads were washed with buffer A containing 20 mM imidazole and eluted with buffer A containing 500 mM imidazole. The tag was cleaved by the TEV protease during a 2 hrs dialysis at 4°C against 50 mM Tris-HCl pH 8.0, 150 mM NaCl, 1 mM EDTA 2 mM DTT. The sample was then boiled 10 min at 95 °C, spun down 5 min at 16,000 *xg* to remove thermo-sensitive contaminants and injected on Superdex 75 pg (GE Healthcare) equilibrated with 50 mM HEPES 1 mM EDTA pH 7.0. Sample concentration was calculated using its estimated molecular extinction coefficient of 9,970 M^-1^ cm^-1^ at 280 nm. The protein sample was characterized for folding using NMR HSQC spectra, before and after the heating at 95°C.

For BRCA2_190-284_, clarified cell lysate was loaded onto Glutathione (GSH) Sepharose beads (GE Healthcare) equilibrated with buffer A. After 2 h of incubation at 4°C, beads were washed with buffer A and eluted with buffer A containing 20 mM reduced glutathione. The tag was cleaved by the precision protease during an overnight dialysis at 4°C against buffer B (50 mM HEPES pH 7.0, 1 mM EDTA) with 2 mM DTT and 150 mM NaCl. The cleaved GST-tag was removed by heating the sample for 15 min at 95°C and spun it down for 10 min at 16,000 x *g*. Sample concentration was calculated using its estimated molecular extinction coefficient of 10,363 M^-1^ cm^-1^ at 280 nm. The protein sample was characterized for folding using NMR HSQC spectra, before and after the heating at 95°C. BRCA2_190-284_ was dialyzed overnight at 4°C against buffer B with 2 mM DTT.

**Expression and purification of PLK1 and PLK1-kinase dead (PLK1-KD)**

The recombinant 6xHis-PLK1 and 6xHis-PLK1-K82R mutant (PLK1-KD) were produced in sf9 insect cells by infection for 48h (28°C, 110 rpm shaking) with the recombinant baculovirus (PLK1-pFast-Bac HT vector). Infected cells were collected by centrifugation (1300 rpm, 10 min, 4°C), washed with 1xPBS, resuspended in lysis buffer (1xPBS, 350 mM NaCl, 1% Triton X-100, 10% glycerol, EDTA-free Protease Inhibitor Cocktail (Roche), 30 mM imidazole). After 1h rotation at 4°C the lysate was centrifuged (25000 rpm, 1h, 4°C) and the supernatant was collected, filtered (0.4 µm) and loaded immediately onto a Ni-NTA column (Macherey Nagel) equilibrated with Buffer A1 (1xPBS with 350 mM NaCl, 10% glycerol and 30 mM imidazole, the column was washed with buffer A2 (1xPBS with 10% glycerol) and the protein was eluted with Buffer B1 (1x PBS with 10% glycerol and 250 mM imidazole). The eluted protein was diluted to 50 mM NaCl with Buffer A before being loaded onto a cationic exchange Capto S column (GE Healthcare) equilibrated with Buffer A1cex (50 mM HEPES (pH 7.4), 50 mM NaCl and 10% glycerol), the column was washed with Buffer A1cex before elution with Buffer B1cex (50 mM HEPES (pH 7.4), 2M NaCl and 10% glycerol). The quality of the purified protein was analysed by SDS-PAGE and the proteins concentration was determined using Bradford protocol with BSA as standard. The purest fractions were pooled and dialyzed against storage buffer (50 mM Tris-HCl (pH7.5), 150 mM NaCl, 0.25 mM DTT, 0.1 mM EDTA, 0.1 mM EGTA, 0.1 mM PMSF and 25% Glycerol) and stored in -80°C. The purified proteins can be seen in Supplementary Fig. 10.

**Expression and purification of PLK1_PBD_**

The pT7-6His-Sumo-PLK1 PBD (326-603) plasmid was expressed in Tuner pLacI pRare cells (Protein Expression and Purification Core Facility, Institut Curie), 2L of TB medium with Kanamycin and Chloramphenicol antibiotics were inoculated with cells from the pre-culture. The cells were grown at 37°C until an OD_600_ of ~ 0.85. The temperature was decreased to 20°C and the expression was induced by 1mM IPTG overnight. The cells were harvested by 15 min of centrifugation at 4690 *x g*, at 4 °C. The cell pellets were suspended in 80 ml of 1 x PBS, pH 7.4, 150 mM NaCl, 10% glycerol, EDTA-free Protease Inhibitor Cocktail (Roche), 5 mM β-mercapto-ethanol (β-ME). The suspension was treated with benzonase nuclease and MgCl_2_ at 1 mM final concentration for 20 min at 4°C. The suspension was lysed by disintegration at 2 kbar (Cell distruptor T75, Cell D) followed by centrifugation at 43000 *x g*, for 45 min, at 4 °C. The supernatant was loaded at 1 ml/min on a His-Trap FF-crude 5 mL column (GE Healthcare) equilibrated with PBS buffer, pH 7.4, 150 mM NaCl, 10% glycerol, 5 mM β-ME (A) and 20 mM imidazole. The proteins were eluted in a linear gradient from 0 to 100 % with the same buffer (A) containing 200 mM imidazole, over 10 column volumes (CV). The purest fractions were pooled and dialyzed (8 kDa cut-off) against 20 mM Tris-HCl buffer, pH 8.0, 100 mM NaCl, 0.5 mM EDTA, 5 mM β-ME, 10% glycerol at 4 °C. 6xHis-SUMO Protease (Protein Expression and Purification Core Facility, Institut Curie) was added at 1/100 (w/w) and incubated overnight at 4 °C to cleave the 6His-SUMO tag. The cleaved PBD-PLK1 was purified using Ni-NTA agarose resin (Macherey Nagel), washed with the following buffer: 20 mM Tris-HCl pH 8.0, 100 mM NaCl, 0.5 mM EDTA, 5mM β-ME and 10 % glycerol. The sample was incubated with the resin for 1h at 4 °C and the flow-through was collected. The sample was concentrated on an Amicon Ultra Centrifugal Filter Unit (10 kDa cut-off) and injected at 0.5 ml/min on a Hi-Load 16/60 Superdex column (GE healthcare), equilibrated with 20 mM Tris-HCl buffer, pH 8.0, 100 mM NaCl, 0.5 mM EDTA, 5 mM β-ME. The protein concentration was estimated by spectrophotometric measurement of absorbance at 280 nm. The purified protein is shown in Supplementary Fig. 10.

The GST-tagged PLK1_PBD_ (365-603) was expressed in *E. coli* BL21 (DE3) STAR cells, induced with 0.5 mM IPTG at an OD_600_ of 0.6, and grown at 37 °C for 3h. The PBD (365-603) was purified by glutathione affinity chromatography. After GST cleavage (using a 6His-TEV protease), the tag and the protease were retained using GST- and NiNTA-agarose affinity chromatography, and the PBD collected in the flow-through was further purified by gel filtration chromatography. The protein was dialyzed against a buffer containing 50 mM Tris-HCl pH 8, NaCl 150 mM, and 5 mM β-ME.

***In vitro* PLK1 kinase assay**

0.5 µg purified 2xMBP-BRCA2_1-250_ or 25 ng RAD51 protein, was incubated with recombinant active PLK1 (0, 50 or 100 ng) or PLK1-kinase dead (100 ng) (purchased from Abcam or purified from sf9 insect cells as detailed above, see Figure EV11B for the comparison of the kinase activity of both PLK1 preparations) in kinase buffer (25 mM HEPES, pH 7.6, 25 mM ß-glycerophosphate, 10 mM MgCl_2_, 2 mM EDTA, 2mM EGTA, 1 mM DTT, 1 mM Na_3_VO_4_, 10 µM ATP and 1 µCi [γ^32^P] ATP (Perkin Elmer)) in a 25 µl total reaction volume. After 30 min incubation at 30°C the reaction was stopped by heating at 95°C for 5 min in SDS-PAGE sample loading buffer. The samples were resolved by 7.5 % SDS-PAGE and [γ^32^P] ATP labelled bands were analysed with PhosphorImager (Amersham Bioscience) using ImageQuant^TM^ TL software (GE Healthcare Life Science). To control for the amount of substrate in the kinase reaction, before adding [γ^32^P] ATP, half of the reaction was loaded on a 7.5 % stain free SDS-PAGE gel (BioRad), the protein bands were visualized with ChemiDoc XRS+ System (BioRad) and quantified by Image Lab^TM^ 5.2.1 Software (BioRad). The relative phosphorylation of 2xMBP-BRCA2_1-250_ was quantified as ^32^P-labelled 2xMBP-BRCA2_1-250_ (ImageQuant^TM^ TL software) divided by the intensity of the 2xMBP-BRCA2_1-250_ band in the SDS-PAGE gel (Image Lab^TM^ 5.2.1 Software). In the control experiment where PLK1 inhibitor was used, 50 nM BI2536 (Selleck Chemicals) was added to the kinase buffer.

***In vitro* protein binding assay**

To assess the interaction between recombinant PLK1 and BRCA2_1-250_ after phosphorylation by PLK1, a kinase assay was performed with 0.2 µg recombinant PLK1 or PLK1-kinase dead (PLK1-KD) and 0.5 µg purified 2xMBP-BRCA2_1-250_ (WT or the VUS T207) in kinase buffer supplemented with 250 µM ATP (no [γ^32^P] ATP) in a total reaction volume of 20 µl, one control reaction without ATP was performed with PLK1 and 2xMBP-BRCA2_1-250_ WT. After 30 minutes incubation at 30°C, 15 µl amylose beads was added to the reaction and incubated for 1h at 4°C. The beads were centrifuged at 2000 x g for 2 minutes at 4°C and the unbound fraction was collected before the beads were washed three time in kinase buffer (no ATP) containing 0.5% NP-40 and 0.1% Triton X-100. Bound proteins were eluted from the beads with 10 mM maltose, protein complexes were separated by SDS-PAGE and analysed by western blotting. To control for the amount of proteins in the reaction, 2 µl of the kinase reaction (before adding the amylose beads) was loaded as input. The protein bands were visualized with ChemiDoc XRS+ System (BioRad) and quantified by Image Lab^TM^ 5.2.1 Software (BioRad). The relative pull-down of PLK1 was quantified as the intensity of the PLK1 band in the pull-down divided by the intensity of the PLK1 band in the input (ImageQuant^TM^ TL software).

To discard possible remaining phosphorylation of the 2xMBP-BRCA2_1-250_ fragment coming from the purification of the protein from HEK293T cells, the 2xMBP-BRCA2_1-250_ fragment (WT or the variant T207) was incubated with kinase buffer (no added ATP) supplemented with FastAP Thermosensitive Alkaline Phosphatase (Thermo Fisher Scientific Cat. #EF0654) for 1h at 37°C before addition of recombinant PLK1 followed by 30 min incubation at 30°C and 1h incubation with amylose beads as described above.

**NMR spectroscopy**

NMR experiments were carried out at 283 K on 600 and 700 MHz Bruker spectrometers equipped with a triple resonance cryoprobe. For NMR signal assignments, standard 3D triple resonance NMR experiments were recorded on ^15^N and ^13^C labelled samples of BRCA2_48-218_ WT and BRCA2_190-284_ WT and T207A. Analyses of these experiments provided backbone resonance assignment for the non-phosphorylated and phosphorylated forms of these BRCA2 fragments. To follow the PLK1 phosphorylation kinetics, the ^15^N labelled fragment BRCA2_48-218_ (50 µM) was mixed to a first PLK1 sample at 0.1 µM and the ^15^N labelled fragment BRCA2_190-284_ (200 µM) was mixed to another PLK1 sample at 1.1 µM. The mixes were incubated at pH 7.8 and 298 K. For each time point, a 140 µl sample was heated during 10 min at 368 K to inactivate PLK1, D_2_O was added, the pH was adjusted to 7.0 and a ^1^H-^15^N SOFAST-HSQC experiment ^51^ was recorded. The HSQC experiments were performed using 2048 X 256 time points, 64 scans and an interscan delay of 80 ms. Data processing and analysis were carried out using Topspin and CcpNmr Analysis 2.4.2 softwares.

**Analysis of phosphorylation assays followed by NMR**

In the HSQC spectra, the intensity of peaks of the phosphorylated residues pS193 and pT207, as well as the intensity of peaks corresponding to their non-phosphorylated form was retrieved at each time point of the kinetics. In order to estimate the fraction of phosphorylation for each residue at each point, the function Intensity_(phospho)_ = f[Intensity_(non-phospho)_] was drawn for each residue, the trendline was extrapolated to determine the intensity corresponding to the 100% phosphorylated residue and then the percentage of phosphorylation could be calculated at each time point by dividing peak intensities corresponding to the phosphorylated residue by the calculated intensity at 100% phosphorylation. Peaks corresponding to residues closed to a phosphorylated residue (L209 and V211 for pT207; D191, S197 and S195 for pS193) and thus affected by this phosphorylation were also treated using the same protocol and they were used to obtain a final averaged curve of the evolution of the percentage of phosphorylation at positions 193, 207 with time.

**Isothermal Titration Calorimetry**

ITC measurements were performed with the PLK1 PBD protein (amino acid 326 to amino acid 603) and BRCA2 peptides in 50 mM Tris-HCl buffer, pH 8.0 containing 150 mM NaCl and 5 mM β-ME, using a VP-ITC instrument (Malvern), at 293 K. We used automatic injections of 8 or 10 µl. The titration data were analyzed using the program Origin 7.0 (OriginLab) and fitted to a one-site binding model. To evaluate the heat of dilution, control experiments were done with peptide or protein solutions injected into the buffer. The peptides used for the ITC experiments were synthesized by GeneCust (Ellange, LU) or Genscript (Piscataway, NY). The peptides were acetylated and amidated at the N-terminal and C-terminal ends, respectively (see Table 7 for peptide sequences). Only peptide BRCA2_190-284_ was expressed in bacteria and purified as detailed above (see “Expression and purification of BRCA2_190-284_ for NMR” section).

**Crystallization and Structure Determination**

The purified PBD protein (amino acid 365 to amino acid 603) was concentrated to 6 mg/ml, and mixed to the ^194^WSSSLATPPTLSS{pT}VLI^210^ (pT207) BRCA2 peptide at a 3:1 molar ratio. The crystals were obtained by hanging drop vapor diffusion method at room temperature (293 K), by mixing 1µl of complex with 1µl of solution containing 10% PEG 3350, 100 mM BisTris pH 6.5, and 5 mM DTT. Diffraction data were collected at the Proxima 1 beamline (SOLEIL synchrotron, Gif-sur-Yvette, France)*.* The dataset was indexed and integrated using XDS through the autoPROC package^52^. The software performs an anisotropic cut-off (Tickle *et al*., STARANISO (2018) Global Phasing Ltd.) of merged intensity data, a Bayesian estimation of the structure amplitudes, and applies an anisotropic correction to the data. The structure was solved by molecular replacement using PHENIX (Phaser) software^53^. Two molecules of PBD were consecutively positioned. Electron density for the peptide was clearly visible in the position previously reported in other PBD structures in complex with phosphorylated peptides (PDB 4O56 or 3P35). Refinement was performed using BUSTER^54^ and PHENIX^55^. The model was built with Coot^56^. A summary of crystallographic statistics is shown in Supplementary Table 2. The figures were prepared using Pymol v.1.7.4.0 (Schrödinger, LLC).

**Generation of stable DLD1 clones**

For generation of DLD1 BRCA2^-/-^ cell lines stably expressing human BRCA2 variants of interest, we transfected one 100 mm plate of DLD1 BRCA2^-/-^ cells at 70% of confluence with 10 µg of a plasmid containing human EGFP-MBP-tagged BRCA2 cDNA (corresponding to accession number NM_000059) using TurboFect (Thermo Fisher Scientific), 48h post-transfection the cells were serial diluted and cultured in media containing 1 mg/ml G418 (Sigma-Aldrich) for selection. Single cells were isolated and expanded. To verify and select the clones, cells were resuspended in cold lysis buffer H (50 mM HEPES (pH 7.5), 250 mM NaCl, 1% NP-40, 5 mM EDTA, 1 mM DTT, 1 mM PMSF and EDTA-free Protease Inhibitor Cocktail (Roche)), incubated on ice for 30 min, sonicated and centrifuged at 10,000 x *g* for 15 min, 100 µg total protein lysate was run on a 4-15% SDS-PAGE followed by immunoblotting using BRCA2 and GFP antibodies to detect EGFP-MBP-BRCA2. Clones with similar expression levels were selected for functional studies.

The presence of the point mutations in the genome of the clones was confirmed by extraction of genomic DNA using Quick-DNA^TM^ Universal Kit (ZYMO Research) followed by amplification of the N-terminal of BRCA2 (aa 1-267) by PCR using a forward primers that binds to the end of MBP and a reverse primer that binds to amino acid 267 in BRCA2, the presence of the point mutations was confirmed by sequencing of the PCR product (see Table 2 and Table 8 for primer sequences).

**Cell extracts, immunoprecipitation and western blotting**

For the interaction between BRCA2_1-250_ and endogenous PLK1, U2OS cells were transfected with 2xMBP-BRCA2_1-250_ construct (WT, M192T, S193A, S196N, T200K, S206C, and T207A) using TurboFect (Thermo Fisher Scientific), 30 h post-transfection cells were synchronized by nocodazole (300 ng/ml), harvested and lysed in extraction buffer A (20 mM HEPES (pH 7.5), 150 mM NaCl, 0.1% NP40, 2 mM EGTA, 1.5 mM MgCl_2,_ 50 mM NaF, 10 % glycerol, 1 mM Na_3_VO_4_, 20 mM ß-glycerophosphate, 1 mM DTT and EDTA-free Protease Inhibitor Cocktail (Roche)). After centrifugation at 18,000 x *g* for 15 min, the supernatant was incubated with amylose resin (NEB) for 1.5h at 4°C. The beads were washed five times in extraction buffer before elution with 10 mM maltose. Bound proteins were separated by SDS-PAGE and analysed by western blotting. Where PLK1 and CDK1 inhibitor was used, the cells were synchronized in mitosis by nocodazole (14h) followed by 2h treatment with PLK1 inhibitor (50-100 nM BI2536 (Selleck Chemicals) or 50 µM BTO-1 (Sigma-Aldrich)) or the CDK1 inhibitor (10 µM, Ro-3306, (Selleck Chemicals)) before being harvested. The cells were lysed in extraction buffer, pre-cleared by centrifugation and total protein lysate was separated by SDS-PAGE and analysed by western blotting. Where proteasome inhibitor was used during the mitotic block, the cells were synchronized by nocodazole for 14h before the MG-132 (50 µM, Sigma-Aldrich) was added to the media and the cells were cultured for additional 2h before harvesting.

For analysis of BUBR1 and pBUBR1 levels in mitosis, nocodazole (100 ng/ml) treated DLD1 BRCA2^-/-^ clones were lysed in extraction buffer A, pre-cleared by centrifugation and total protein lysate was separated by SDS-PAGE and analysed by western blotting.

For analysis of the interaction between BRCA2-PLK1, BRCA2-BUBR1 and for the protein complex BRCA2-pBUBR1/BUBR1-PP2A(C)-PLK1 in mitosis, DLD1 BRCA2^-/-^ stable clones expressing EGFP-MBP-BRCA2 (WT or the VUS S206C or T207A) were synchronized with nocodazole, harvested and lysed in extraction buffer A. The lysate were pre-cleared by centrifugation before incubation with GFP-TRAP beads (Chromotek) for 2h at 4°C to pull-down EGFP-MBP-BRCA2. Around 3 mg total protein lysate was used per pull-down. The beads were washed 5 times in extraction buffer and 2 times in extraction buffer with 500 mM NaCl. Bound proteins were eluted by boiling the samples for 4 min in 3x SDS-PAGE sample loading buffer (SB), eluted proteins were separated by SDS-PAGE and analysed by western blotting using anti-mouse PLK1, anti-mouse BUBR1, anti-rabbit pT680-BUBR1, anti-mouse PP2A-C and anti-mouse BRCA2 (OP95) antibodies.

For immunoprecipitation of endogenous BUBR1, nocodazole treated DLD1 BRCA2^-/-^ stable clones expressing BRCA2 WT or the variants (S206C or T207A) were lysed in extraction buffer A. After centrifugation, 2000-3000 µg total protein lysate was pre-cleared by incubation with 20 µl Protein G PLUS-Agarose (Santa Cruz, sc-2002) for 30 min at 4°C. The pre-cleared lysate was incubated with 1.25 µg BUBR1 mouse antibody or control mouse IgG over night at 4°C before addition of 40 µl Protein G PLUS-Agarose, the lysate was incubated for additional 30 min before immunoprecipitates were collected by centrifugation. After four washes in extraction buffer and two washes in extraction buffer with 500 mM NaCl, the beads were re-suspended in SB, boiled and the immunocomplexes were analysed by western blotting using anti-rabbit BUBR1, anti-mouse PLK1 and anti-mouse PP2A-C antibodies.

For the interaction between the phosphomimic BUBR1-3D mutant (S670D, S676D and T680D) and endogenous PP2A, the DLD1 BRCA2^-/-^ stable clones expressing BRCA2 WT or the S206C variant was transient transfected with the pcDNA3-3xFLAG-BUBR1-3D-RFP construct. The transfection media was replaced 30h post-transfection with fresh growth media containing 0.1µg/ml nocadozole and the cells were incubated additional 14h before harvesting. The cells were lysed and an immunoprecipitation was performed as described above for the BUBR1 immunoprecipitation using rabbit anti-tRFP antibody (Cat.# AB233, Evrogen) to pull-down the 3xFLAG-BUBR1-3D-RFP protein. Immunocomplexes were analysed by western blotting using anti-mouse BUBR1 and anti-mouse PP2A-C antibodies.

For all Western blots, the protein bands were visualized with ChemiDoc XRS+ System (BioRad) and quantified by Image Lab^TM^ 5.2.1 Software (BioRad). For the relative expression levels (Fig. 5g-5i, Supplementary Fig. 5d), the intensity of the band of interest was divided by the intensity of the signal from the stain free gel. The results are presented as percentage compared to BRCA2 WT clone. To calculate the relative co-immunoprecipitation (co-IP)/co-pull-down of a protein of interest, the intensity of the band in the co-IP was divided by the intensity of the band in the input (ImageQuant^TM^ TL software), the ratio co-IP:input of the protein of interest was then divided by the intensity of the band of the immunoprecipitated protein. StainFree images of the gels before transfer were used as loading control for the input and cropped image is shown in the figures.

**Antibodies used for western blotting**

mouse anti-MBP (1:5000, R29, Cat. #MA5-14122, Thermo Fisher Scientific), mouse anti-BRCA2 (1:1000, OP95, EMD Millipore), rabbit anti-GFP (1:5000, Protein Expression and Purification Core Facility, Institut Curie), mouse anti-PLK1 (1:5000, clone 35-206, Cat. #05-844, EMD Millipore), mouse anti-BUBR1 (1:1000, Cat. #612502, BD Transduction Laboratories), rabbit anti-BUBR1 (1:2000, Cat. #A300-386A, Bethyl Laboratories), mouse anti-PP2A C subunit (1:1000, clone 1D6, Cat. #05-421, EMD Millipore), rabbit anti-pT680-BUBR1 (1:1000, EPR 19958, Cat. #ab200061, Abcam), and rabbit anti-pS676-BUBR1 (1:1000, R193, kind gift from Dr. Erich A. Nigg). Horseradish peroxidase (HRP) conjugated 2^nd^ antibodies used: mouse-IgGκ BP-HRP (IB: 1:10 000, Cat. #sc-516102, Santa Cruz), goat anti-rabbit IgG-HRP (IB: 1:5000, Cat. #sc-2054, Santa Cruz), goat anti-mouse IgG-HRP (1:10 000, Cat.# 115-035-003, Interchim), goat anti-rabbit IgG-HRP (1:10 000, Interchim, Cat.# 111-035-003).

**siRNA transfection**

For analysis of the pT680-BUBR1 levels in U2OS cells after transient depletion of endogenous BRCA2 with RNAi, U2OS cells were transfected with 200 nM siRNA targeting the 3’UTR of BRCA2 (siBRCA2 #1: SI00000966, Qiagen) using jetPRIME (Polyplus Transfection, Cat.# 114-07). As control, the cells were transfected with the 200 nM of the si-control RNA (siRNA control ON-TARGETplus Non-targeting Pool. D-001810-10-05, Thermo Scientific) The transfection media was replaced 30h post-transfection with fresh growth media containing 0.1µg/ml nocadozole and the cells were incubated additional 14h before harvesting. Total protein lysate was extracted as described above and 30-50 µg total protein lysate was resolved on a 4-15% SDS-PAGE and analysed by western blotting using anti-mouse BRCA2 (OP95), anti-mouse BUBR1, anti-rabbit pT680-BUBR1 and anti-mouse PLK1 antibodies (see above for reference number).

For depletion of endogenous BRCA2 in DLD1 BRCA2^+/+^ cells for analysis of γH2AX and RAD51 foci, the cells were transfected with a combination of the BRCA2 5’UTR siRNA (SI00000966, Qiagen) and IAC204 (Dharmacon D-003462-04) (100 nM each) or the ON-TARGET plus Non-targeting oligonucleotide D-001810-04-20, Thermo Scientific 100 nM) and fixed and expose to IR 30h post-transfection (see Supplementary Fig. 8c for the Western blot showing BRCA2 depletion) for analysis of DNA repair foci (see Supplementary Fig. 8c for the Western blot showing BRCA2 depletion).

**Phosphatase treatment**

DLD1 BRCA2^-/-^ cells stably expressing EGFP-MBP-BRCA2 WT were synchronized in mitosis by nocodazole (14h), harvested, lysed in extraction buffer A without phosphatase inhibitors (NaF, Na_3_VO_4_ and ß-glycerophosphate), and pre-cleared by centrifugation. Increased amount (0-20U) of FastAP Thermosensitive Alkaline Phosphatase (Thermo Fisher Scientific Cat. #EF0654) was added to 15 µg of total protein lysate in FastAP Buffer in a total reaction volume of 60 µl. After 1h incubation at 37°C the reaction was stopped by heating at 95°C for 5 min in SDS-PAGE sample loading buffer, 30 µl of the reaction was loaded on a 4-15 % SDS-PAGE gel, the gel was transferred onto nitrocellulose membrane and the levels of pT680-BUBR1 were analysed by western blotting.

**Cell survival and viability assays**

For clonogenic survival assay, DLD1 BRCA2^-/-^ cells stably expressing full-length EGFP-MBP-BRCA2 and the variants (S206C and T207A) were treated at 70% of confluence with Mitomycin C (Sigma-Aldrich) at concentrations: 0, 0.5, 1.0 and 2.5 µM. After 1 h drug treatment the cells were serial diluted in normal growth media containing penicillin/streptomycin (Eurobio) and seeded in triplicates into 6-well plates. The media was changed every third day, after 10-12 days in culture the plates were stained with crystal violet, colonies were counted and the surviving fraction was determined for each drug concentration.

Cell viability was assessed with 3-[4,5-Dimethylthiazol-2-yl]-2,5-diphenyltetrazolium bromide (MTT, #M5655, Sigma Aldrich) after treatment with MMC and the PARP inhibitor Olaparib (AZD2281, Ku-0059436, #S1060, Selleck Chemicals). For MMC, the cells were plated in triplicates in 96-well microplates (3000-5000 cells/well) the day before treatment. The cells were washed once in PBS before addition of serum-free media containing MMC at the concentrations: 0, 1.0 and 2.5 µM. After 1h treatment the cells were washed once in PBS and incubated for 72h in normal growth media before the viability was measured by MTT assay. For PARP inhibition, the cells were seeded 4h before 4-days treatment in normal growth media with Olaparib at concentrations: 0, 2.5 and 5.0 µM.

**Homologous recombination assays**

We applied a DSB-mediated gene targeting strategy using site-specific TALEN nucleases to quantify HR in cells. DLD1 BRCA2^-/-^ cells stably expressing full-length GFPMBP-BRCA2 and the variants (S206C and T207A) were transfected using AMAXA technology (Lonza) nucleofector kit V (Cat. # VCA-1003) with 3 µg of the promoter-less donor plasmid (AAVS1-2A-mCherry) with or without 1 µg of each AAVS1-TALEN encoding plasmids (TALEN-AAVS1-5’ and TALEN-AAVS1-3’, see Table 10 for sequences, kind gift from Dr. Carine Giovannangeli). For each transfection, 1 x 10^6^ cells were transfected using program L-024, the cells were seeded in 6-well plate in culture media without selection antibiotics. The day after transfection the media was changed to media with selection and 48h post-transfection the cells were trypsinized and reseeded on a 10-cm culture dish and cultured for additional five days. The percentage of mCherry positive cells was analysed on a BD FACSAria III (BD Bioscience) using FACSDiva software and data was analysed with FlowJo 10.4.2 software (Tree Star Inc.).

**Analysis of tetraploid cells**

For the analysis of S-phase tetraploid cells in the DLD1 BRCA2^-/-^ stable clones, the cells were incubated with 10 µM BrdU for 20 minutes before they were harvest, fixed and stained for cell cycle analysis using a APC-BrdU flow kit (BD Bioscience, Cat. #552598) following the manufacturer’s instructions.

Labelled cells were analysed on a BD FACSCanto II (BD Bioscience) using FACSDiva software and data were analysed with FlowJo 10.4.2 software (Tree Star Inc.).

**Immunofluorescence**

*Kinetochore localization*

For staining of pT680-BUBR1 and PLK1 at the kinetochore, DLD1 BRCA2^-/-^ stable clones expressing EGFP-MBP-BRCA2-WT or the variant T207A were seeded on coverslips and treated with nocodazole (0.25 µg/ml) for 4h, fixed with 4% PFA in PBS containing 0.5% Triton X-100 for 20 minutes at room temperature. The coverslips were rinsed three times in PBS-T and blocked for 30 minutes with 4% BSA in PBS before incubation with primary antibodies (human anti-CREST (1:100, Cat. #15-234-0001, Antibodies Online) together with either rabbit anti-pT680-BUBR1 (1:500, clone EPR 19958, Abcam, Cat. #ab200061) or mouse anti-PLK1 (1:500, clone F-8, Santa Cruz Biotechnology, Cat. #sc-17783), diluted in PBS-T with 5% BSA over night at 4°C. After three washes of 5 minutes in PBS-T the coverslips were incubated for 2h incubation at room temperature with respective Alexa Fluor conjugated secondary antibody (for pBUBR1; goat anti-human Alexa-488 (1:1000, Cat. # A11013, Life Technologies) and donkey anti-rabbit Alexa-488 (1:1000, Cat. #A-21206, Thermo Fisher Scientific), for PLK1; goat anti-human Alexa-633 (1:500, Cat. # A21091, Life Technologies) together with either donkey anti-rabbit Alexa-488 (1:1000, Cat. #A-21206, Thermo Fisher Scientific) or donkey anti-mouse Alexa-488 (1:1000, Cat. #A-21202, Thermo Fisher Scientific) diluted in PBS-T with 5% BSA. After two washes of 5 min in PBS-T and one rinse in PBS the coverslips were mounted on microscope slides.

For staining of BRCA2 at the kinetochore, U2OS was transient transfected with GFPMBP-BRCA2 construct using TurboFect (Thermo Fisher Scientific). The cells were seeded on coverslips 24h post-transfection and incubated for another 24h before nocadozole (0.25 µg/ml) was added. The cells were treated for 4h followed by fixation as described above for kinetochore localization of pBUBR1, pAuroraB and PLK1. BRCA2 was detected by rabbit anti-BRCA2 (1:500, CA1033, EMD Millipore) and Alexa-488 secondary antibody (donkey anti-rabbit Alexa-488 (1:1000, Cat. #A-21206, Thermo Fisher Scientific), CREST was detected by human anti-CREST (1:100, Cat. #15-234-0001, Antibodies Online) and Alexa-633 secondary antibody (1:500, Cat. # A21091, Life Technologies)), diluted in PBS-T with 5% BSA.

Phosphatase inhibitors (50 mM NaF, 1 mM Na_3_VO_4_, and 20 mM ß-glycerophosphate) were added to all buffers.

*Chromosome alignment and segregation*

DLD1 BRCA2^-/-^ cells and the stable clones expressing EGFP-MBP-BRCA2 WT or the variants (S206C and T207A) were seeded on coverslips in 6-well tissue culture plates and synchronized in mitosis. For analysis of chromosome alignment, the cells were synchronized by double thymidine (2.5 mM, Sigma-Aldrich) block, released for 9h followed by treatment with Monastrol (100 µM, Sigma-Aldrich) for 16h. After incubation with Monastrol the cells were washed twice in PBS before 1h incubation in media containing the proteasome inhibitor MG-132 (10 µM, Sigma-Aldrich). For chromosome segregation analysis, the cells were synchronized by double thymidine block and released in normal growth media for 11h.

After synchronization, the cells were fixed with 100% methanol for 15 min at -20ºC, rinsed once in PBS before permeabilization with PBS containing 0.1% Triton-X for 15 min at room temperature. Nonspecific epitope binding was blocked with 4% BSA (Sigma-Aldrich) in PBS. The coverslips were rinsed in PBS, incubated with primary antibody (mouse anti-α-tubulin (1:5000, GT114, Cat. #GTX628802, Euromedex) and human anti-CREST (1:100, Cat. #15-234-0001, Antibodies Online)) diluted in PBS containing 0.1% Tween-20 (PBS-T) and 5% BSA for 1h at room temperature. After incubation, the coverslips were washed three times of 5 min in PBS-T before being incubated for 1h at room temperature with Alexa Fluor conjugated secondary antibody (donkey anti-mouse Alexa-594 (1:1000, Cat. #A-21203, Thermo Fisher Scientific) and goat anti-human Alexa-488 (1:1000, Cat. # A11013, Life Technologies)) diluted in PBS-T with 5% BSA. The coverslips were washed two times of 5 min each in PBS-T followed by one rinse in PBS before being mounted on microscope slides.

For the analysis of chromosome alignment in cells expressing the phosphomimic BUBR1-3D mutant (S670D, S676D and T680D), the DLD1 BRCA2^-/-^ stable clones expressing the T207A variant was transient transfected with the pcDNA3-3xFLAG-BUBR1-3D-RFP construct using TurboFect. The day after transfection the cells were seeded on coverslips in 6-well tissue culture plates, synchronized in mitosis and prepared for immunofluorescence.

*Aneuploidy*

For aneuploidy analysis the cells were treated with nocodazole for 14h (0.1 µg/ml) to enable chromosome spread; the cells were rinsed in PBS, incubated for 10 min with KCl (50 mM) at room temperature before they were spread on coverslips at 900 rpm for 5 min in a Cytospin 4 (Thermo Scientific). The cells were fixed with 3% paraformaldehyde (PFA) in PBS for 20 min followed by 15 min permeabilization in PBS containing 0.1% Triton X-100. The coverslips were rinsed three times in PBS, blocked with 5% BSA in PBS before incubation with human anti-CREST primary antibody (1:100, Cat. #15-234-0001, Antibodies Online) diluted in PBS over night at 4°C. After incubation the coverslips were washed three times of 5 min in PBS before 1h incubation at room temperature with Alexa Fluor conjugated secondary antibody (goat anti-human Alexa-555 (1:1000, Cat. #A-21433, Thermo Fisher Scientific)) diluted in PBS. After three washes of 5 min in PBS the coverslips were mounted on microscope slides.

γ*H2AX and RAD51 foci*

For the detection of γH2AX and RAD51 foci, the cells were seeded on coverslips the day before 6 Gy γ−irradiation (GSR D1, Cs-137 irradiator). Two hours after irradiation, the coverslips were washed twice in PBS followed by one wash in CSK Buffer (10 mM PIPES, pH 6.8, 0.1 M NaCl, 0.3 M sucrose, 3 mM MgCl_2_, EDTA-free Protease Inhibitor Cocktail (Roche)). The cells were permeabilized for 5 minutes at room temperature in CSK buffer containing 0.5% Triton X-100 (CSK-T) followed by one rinse in CSK buffer and one rinse in PBS before fixation for 20 minutes at room temperature with 2% PFA in PBS. After one rinse in PBS and one in PBS-T, the cells were blocked for 5 minutes at room temperature with 5% BSA in PBS-T before incubation for 2h at room temperature with primary antibodies diluted in PBS-T with 5% BSA. After primary antibody incubation, the coverslips were rinsed in PBS-T followed by two washes of 10 minutes in PBS-T and blocked for 5 minutes at room temperature with 5% BSA in PBS-T before incubation for 1h at room temperature with respective Alexa Fluor conjugated secondary antibody diluted in PBS-T with 5% BSA. After one rinse in PBS-T and two washes of 10 minutes in PBS-T the coverslips were rinsed in PBS before being mounted on microscope slides. γH2AX foci were detected by mouse anti-pSer139- γH2AX (1:1000, clone JBW301, EMD-Millipore, Cat. #05-636) and secondary antibody donkey anti-mouse Alexa-594 (1:1000, Cat. #A-21203, Thermo Fisher Scientific). RAD51 foci were detected by rabbit anti-RAD51 (1:100, clone H-92, Santa Cruz Biotechnology, Cat. #sc-8349), followed by secondary antibody donkey anti-rabbit Alexa-488 (1:1000, Cat. #A-21206, Thermo Fisher Scientific).

For the analysis of γH2AX and RAD51 foci in DLD1 BRCA2^+/+^ cells depleted of BRCA2 by siRNA, the DLD1 BRCA2^+/+^ cells were transient transfected with siRNA targeting BRCA2 (see section siRNA transfection above). The day after transfection the cells were seeded on coverslips in 6-well tissue culture plates and radiated as described above.

*Micronuclei*

For analysis of micronuclei, DLD1 BRCA2^-/-^ cells and the stable clones expressing EGFP-MBP-BRCA2 WT or the variants (S206C and T207A) were seeded on coverslips in 6-well tissue culture plates the day before fixation. Centromeres were detected by human anti-CREST primary antibody (1:100, Cat. #15-234-0001, Antibodies Online) and Alexa Fluor conjugated secondary antibody (goat anti-human Alexa-555 (1:1000, Cat. #A-21433, Thermo Fisher Scientific)).

All coverslips were mounted on microscope slides with ProLong Diamond Antifade Mountant with DAPI (Cat. #P36966, Thermo Fisher Scientific).

*Image acquisition and analysis*

For analysis of DNA repair foci, chromosome alignment and segregation, images were acquired in an upright Leica DM6000B wide-field microscope equipped with a Leica Plan Apo 63x NA 1.4 oil immersion objective. The camera used is a Hamamatsu Flash 4.0 sCMOS controlled with MetaMorph2.1 software (Molecular Devices). For Fig. 5J and 6A, 7 to 20 Z-stacks were taken at 0.2 μm intervals to generate a maximal intensity projection image using ImageJ. For the analysis of γH2AX and RAD51 foci, 26 Z-stacks were taken at 0.2 μm intervals to generate a maximal intensity projection using Image J. For the BUBR1-3D-RFP chromosome alignment experiment, RFP negative cells on the same coverslips were used as control.

The number of γH2AX foci per nucleus were counted by a customized macro using a semi-automated procedure; the nucleus was defined by an auto-threshold (Otsu, Image J) on DAPI, a mask was generated and applied onto the Z-projection to count foci within the nucleus. For the definition of foci we applied the threshold plugin IsoData (ImageJ) and for the quantification of foci we used the tool Analyze Particles (ImageJ) setting a range of 5-100 pixels^2^ to select only particles that correspond to the size of a focus. RAD51 foci were quantified using the plugin Find Maxima onto the Z-projection with a prominence of 1000.

For analysis of aneuploidy, kinetochore localization and micronuclei images were acquired in an inverted confocal Leica SP5 microscope with a plan Apo 63x NA 1.4 oil immersion objective with the lasers 405, 488, 561 and 633 nm. For Fig. 7B (aneuploidy), Z-stacks were taken at 0.13 μm intervals to generate a maximal intensity projection image using ImageJ. For the counting of chromosomes in the aneuploidy experiment, the quantification was performed in zoomed areas counting the CREST signal in separated stacks to ensure the counting of all chromosomes. We were able to count up to 65 chromosomes with certainty, thus >65 CREST signals were discarded and not included in the analysis.

For the analysis of kinetochore localization, Z-stacks were taken at 0.21 μm intervals to generate a sum slice projection image using ImageJ, 6 pairs of chromosomes per cell were analysed in 15-21 cells per experiment from two individual experiment. The results from the quantifications (Fig. 5i, Supplementary Fig. 6b) is represented as the ratio between the intensities for pBUBR1/PLK1 and the CREST signal relative to the mean ratio observed for the BRCA2 WT complemented cells. For the images in Fig. 5h (pBUBR1:CREST), Supplementary Fig. 6a (PLK1:CREST) and Supplementary Fig. 8e (micronuclei), Z-stacks were taken to generate a maximal intensity projection image using ImageJ, except for DAPI where the image is from one Z-stack.

**Time-lapse video microscopy of mitotic cells**

For phase-contrast video-microscopy DLD1 BRCA2^-/-^ cells stably expressing full-length EGFP-MBP-BRCA2 and the variants (S206C and T207A) were seeded in 35 mm Ibidi µ-Dishes (Ibidi, Cat. #81156), synchronized by double thymidine block, released and cultured for 4h in normal growth media before the filming was started. The cells were imaged for 16h every 5 min, at oil-40X using an inverted video-microscope (Leica DMI6000) equipped with electron multiplying charge coupled device (EMCCD) camera controlled by Metamorph software (Molecular Devices). Images were mounted using Image J software (1.51s, NIH).

**Statistical analysis**

In all graphs error bars represent the standard deviation (SD) from at least three independent experiments unless otherwise stated, scatter dot plots show median with 95% CI. Statistical significance of differences was calculated with unpaired two-tailed t-test, one/two-way ANOVA with Dunnett’s or Tukey’s multiple comparisons test, Mann-Whitney two-tailed test or Kruskal-Wallis test followed by Dunn’s multiple comparisons test as indicated in the figure legends. All analyses were conducted using GraphPad Prism (version Mac OS X 8.1.2 (227)).

**Materials and Methods tables 1-10**

**Table 1**: Primers used to introduce point mutations in EGFPMBP-BRCA2, 2xMBP-BRCA2_1-250_, GST-BRCA2_190-284_ constructs

| **Mutation** | **Oligo name** | **Sequence (5’-3’)** |
| --- | --- | --- |
| S193A | Fw : oAC543 | CCC ACC CTT AGT TCT GCT GTG CTC ATA GTC |
|  | Rv : oAC544 | GAC TAT GAG CAC AGC AGA ACT AAG GGT GGG |
| M192T | Fw : oAC283 | GTGGATCCTGATACGTCTTGGTCAAGTTC |
|  | Rv : oAC284 | GA ACT TGA CCA AGA CGT ATC AGG ATC CAC |
| S196N | Fw : oAC026 | CCTGATATGTCTTGGTCAAATTCTTTAGCTACACCACC |
|  | Rv : oAC027 | GGTGGTGTAGCTAAAGAATTTGACCAAGACATATCAGG |
| S206C | Fw : oAC028 | CCACCCACCCTTAGTTGTACTGTGCTCATAGTCAG |
|  | Rv : oAC029 | CTGACTATGAGCACAGTACAACTAAGGGTGGGTGG |
| T207A | Fw : oAC545 | GGA TCC TGA TAT GGC TTG GTC AAG TTC TTT AGC |
|  | Rv : oAC546 | GCT AAA GAA CTT GAC CAA GCC ATA TCA GGA TCC |

**Table 2**: Sequencing primers

| **Construct** | **Oligo name** | **Binding site** | **Sequence (5’-3’)** |
| --- | --- | --- | --- |
| GFPMBP-BRCA2, GST-BRCA2_190-284_ | Rv : oAC131 | aa 273 BRCA2 | TTAGTTCGACTTATCCAATGTGGTCTTT |
| 2xMBP-BRCA2_1-250_ | Fw : oAC149 | aa 1-6  BRCA2 | TTATTTGCTAGCCCTATTGGATCCAAAGAG |
| PLK1 | Fw : oAC907 | aa 38 PLK1 | aaagagatcccggaggtcctagtg |

**Table 3:** Primers used to subclone BRCA2_190-284_ into the pGEX-6P-1 vector

| **Construct** | **Oligo name** | **Sequence (5’-3’)** |
| --- | --- | --- |
| BRCA2_190-284_ | Fw: oAC130 | TTAGGATCCATGTCTTGGTCAAGTTCT |
|  | Rv: oAC131 | TTAGTTCGACTTATCCAATGTGGTCTTT |

**Table 4**: Primers used to subclone *PLK1* cDNA into pFastBac HT

| **Primer name** | **Sequence (5’-3’)** |
| --- | --- |
| GA_pFBtev_R | GCCCTGAAAATACAGGTTTTCGGTCGTTGGGAT |
| GA_pFB_UTR_F | TTGTCGAGAAGTACTAGAGGATCATAATCA |
| GA_hPLK_F | ATCCCAACGACCGAAAACCTGTATTTTCAGGGCATGAGTGCTGCAGTGACTGCA |
| GA_hPLK_R | TGATTATGATCCTCTAGTACTTCTCGACAATTAGGAGGCCTTGAGACGGTT |

**Table 5**: Primers used to introduce K82R point mutation in pFastBAC-PLK1 vector to produce PLK1-KD and to subclone PLK1_PBD_ (aa 326-603) into pT7-His6-SUMO

| **Product** | **Oligo name** | **Sequence (5’-3’)** |
| --- | --- | --- |
| K82R-PLK1 | Fw : oAC905 | GCG GGCAGGATTGTGCCTAAG |
|  | Rv : oAC906 | CTTAGGCACAATCCTGCCCGC |
| PLK1_PBD_ | GA_PLKPDBwt_F | ATTGAGGCTCACCGCGAACAGATTGGTGGCTCGATTGCTCCCAGCAGCCT |
|  | GA_PLKPDBwt_R | TTCCTTTCGGGCTTTGTTAGCAGCCGGTCATTAGGAGGCCTTGAGACGGT |

**Table 6**: Primers used to introduce S670D, S676D and T680D point mutations in pcDNA3-3xFlagBUBR1-RFP construct

| **Name** | **Sequence (5’-3’)** |
| --- | --- |
| oAC884 | CAA GAA GCT GGA CCC AAT TAT TGA AGA CGA TCG TGA AGC CGA CCA CTC CTC |
| oAC885 | GAG GAG TGG TCG GCT TCA CGA TCG TCT TCA ATA ATT GGG TCC AGC TTC TTG |

**Table 7**: Synthetic peptide sequences for Isothermal Titration Calorimetry (ITC) and X-ray crystallography

| **Peptide** | **Sequence** |
| --- | --- |
| pS197 | DMSWSS{pS}LAT |
| T207 | WSSSLATPPTLSSTVLI |
| pT207 | WSSSLATPPTLSS{pT}VLI |
| T207A | WSSSLATPPTLSSAVLI |
| T207D | WSSSLATPPTLSSDVLI |
| CpT207 | WSSSLATPPTLSC{pT}VLI |

**Table 8**: Primers for amplifying BRCA2 (aa 1-267) from genomic DNA

| **Primer name** | **Sequence (5’-3’)** |
| --- | --- |
| Fw : OAC035 | GGTCGTCAGACTGTCGATGAAGCC |
| Rv : OAC056 | CAAAGAGAAGCTGCAAGTCATGGATTTGAAAAAACATCAGGG |

**Table 9**: Oligos used for replacing GFP in AAVS1-2A-GFP to mCherry using Gibson Assembly strategy

| **Construct** | **Oligo name** | **Sequence (5’-3’)** |
| --- | --- | --- |
| AAVS1-2A | Fw: oAC537 | TAAAGCGGCCGCGTCGAGTCTAGAGGG |
|  | Rv: oAC538 | CATCTCGAGCCTAGGGCCGGG |
| AAVS1-2A-mCherry | Fw: oAC539 | cccggccctaggctcgagatgGTGAGCAAGGGCGAGGAGGATAAC |
|  | Rv: oAC540 | ctctagactcgacgcggccgctttaCTTGTACAGCTCGTCCATGCCGC |

**Table 10**: TALEN sequences used in DSB-mediated gene targeting assay

| **Name** | **Sequence (5’-3’)** |
| --- | --- |
| TALEN AAVS 5' | TCCCCTCCACCCCACAGT |
| TALEN AAVS 3' | AGGATTGGTGACAGAAAA |
