## Supplementary material for "Proper chromosome alignment depends on BRCA2 phosphorylation by PLK1": Suppl. Figures 1-10

Supplementary Fig. 1

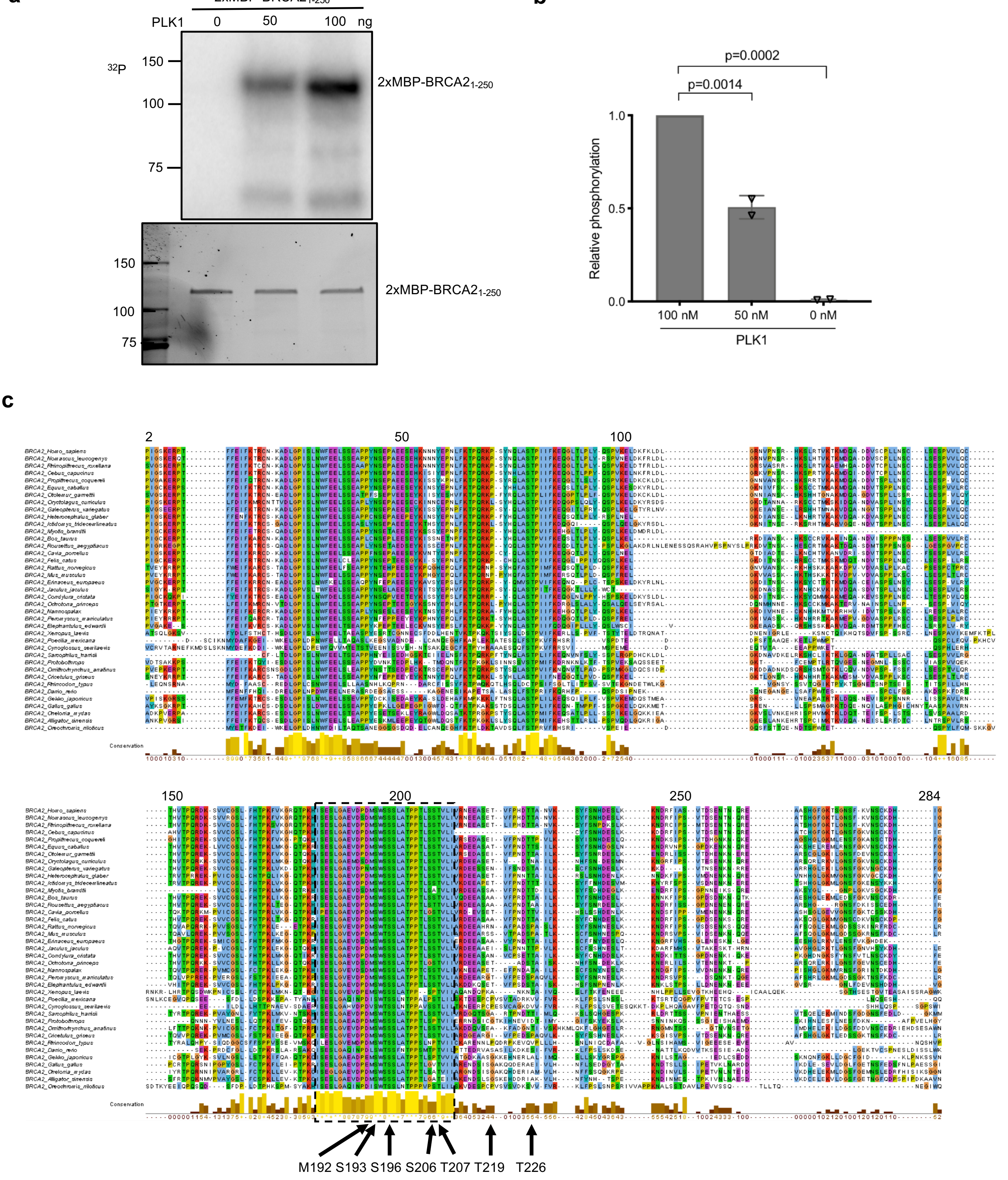

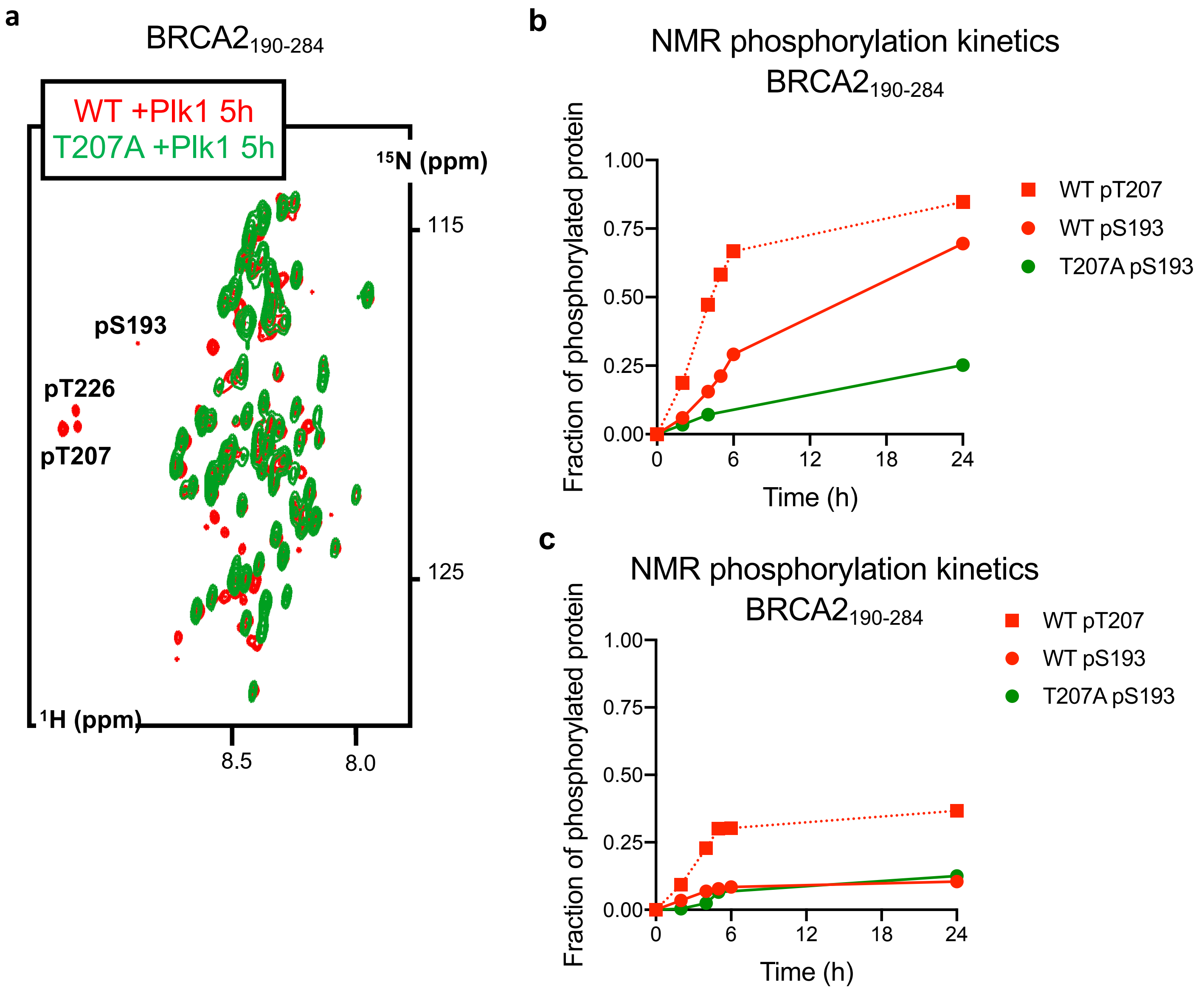
a
BRCA2190-284
WT +Plk1 5h
T207A +Plk1 5h
15N (ppm)
115
125
pS193
pT226
pT207
1H (ppm)
8.5
8.0
b
NMR phosphorylation kinetics
BRCA2190-284
Fraction of phosphorylated protein
1.00
0.75
0.50
0.25
0.00
0
6
12
18
24
Time (h)
WT pT207
WT pS193
T207A pS193
c
NMR phosphorylation kinetics
BRCA2190-284
Fraction of phosphorylated protein
1.00
0.75
0.50
0.25
0.00
0
6
12
18
24
Time (h)
WT pT207
WT pS193
T207A pS193

### Supplementary Fig. 3

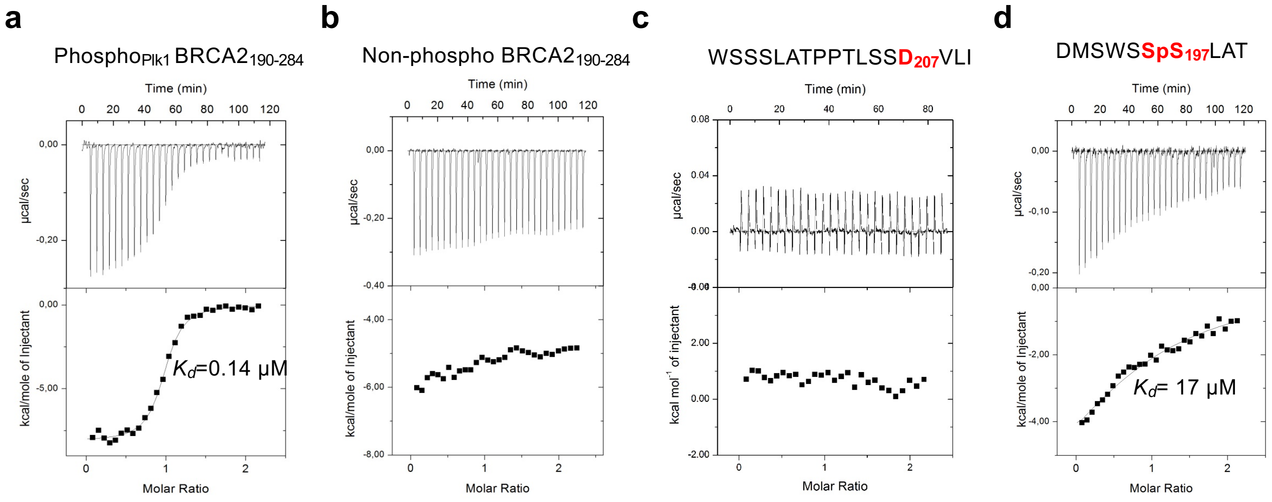

Supplemnetary Fig. 4

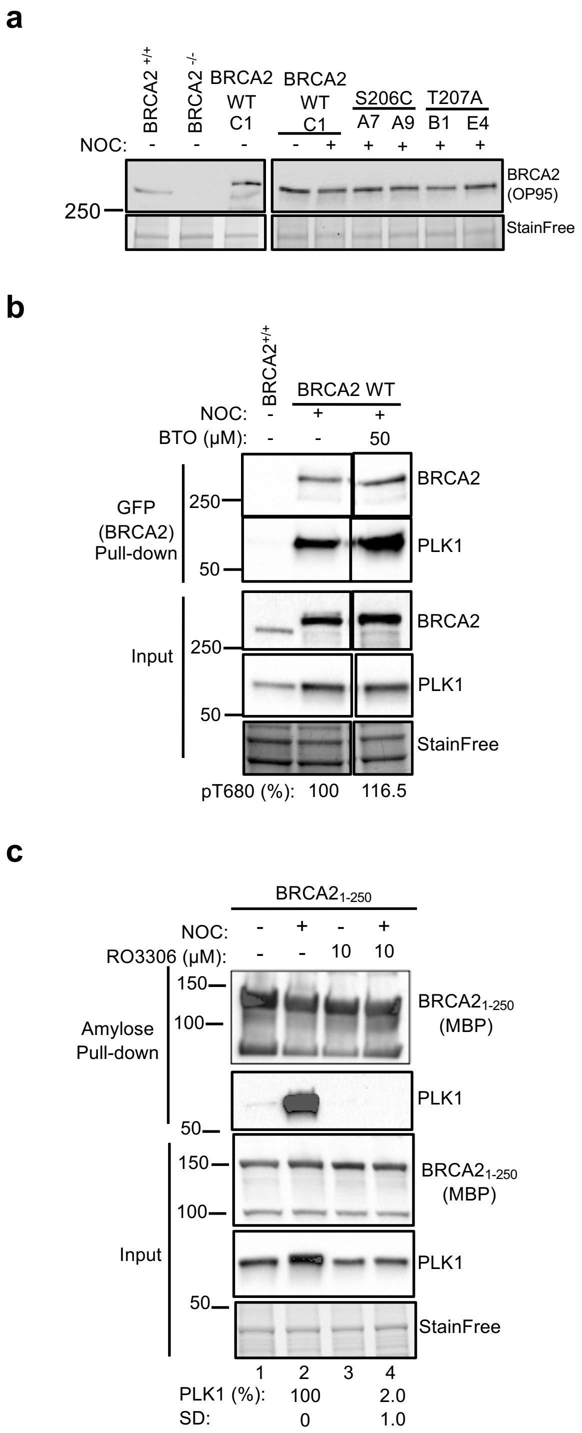

Supplementary Fig 5

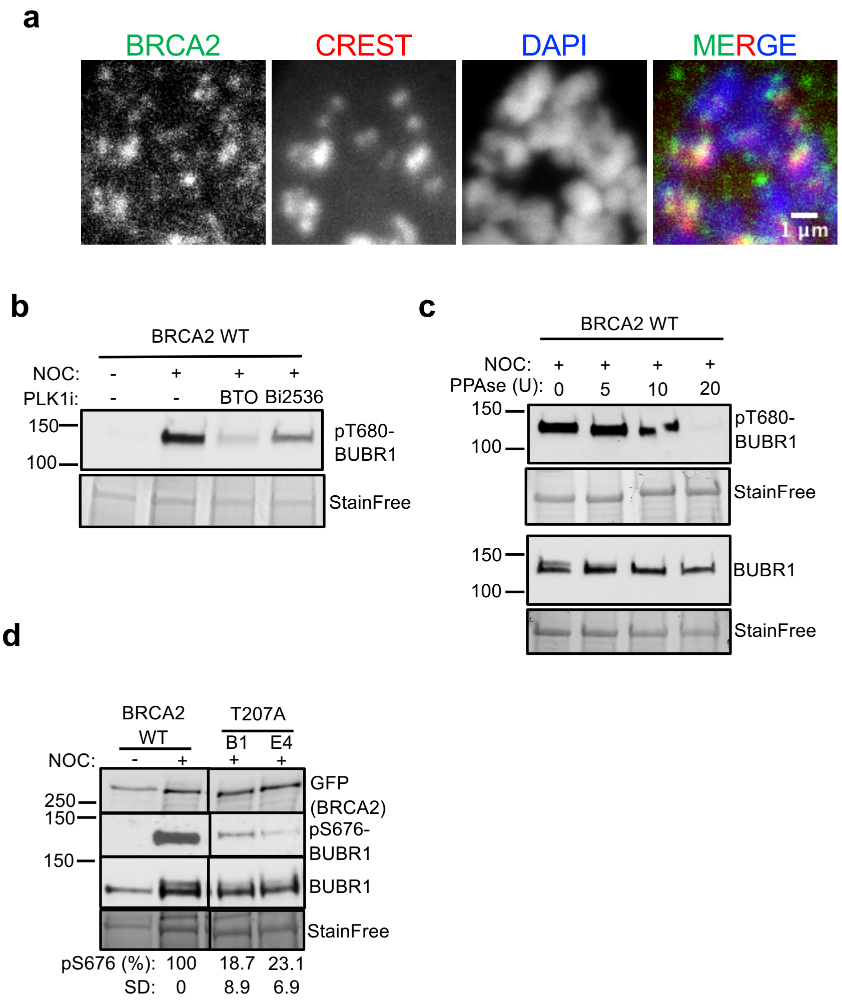

Supplementary Fig. 6

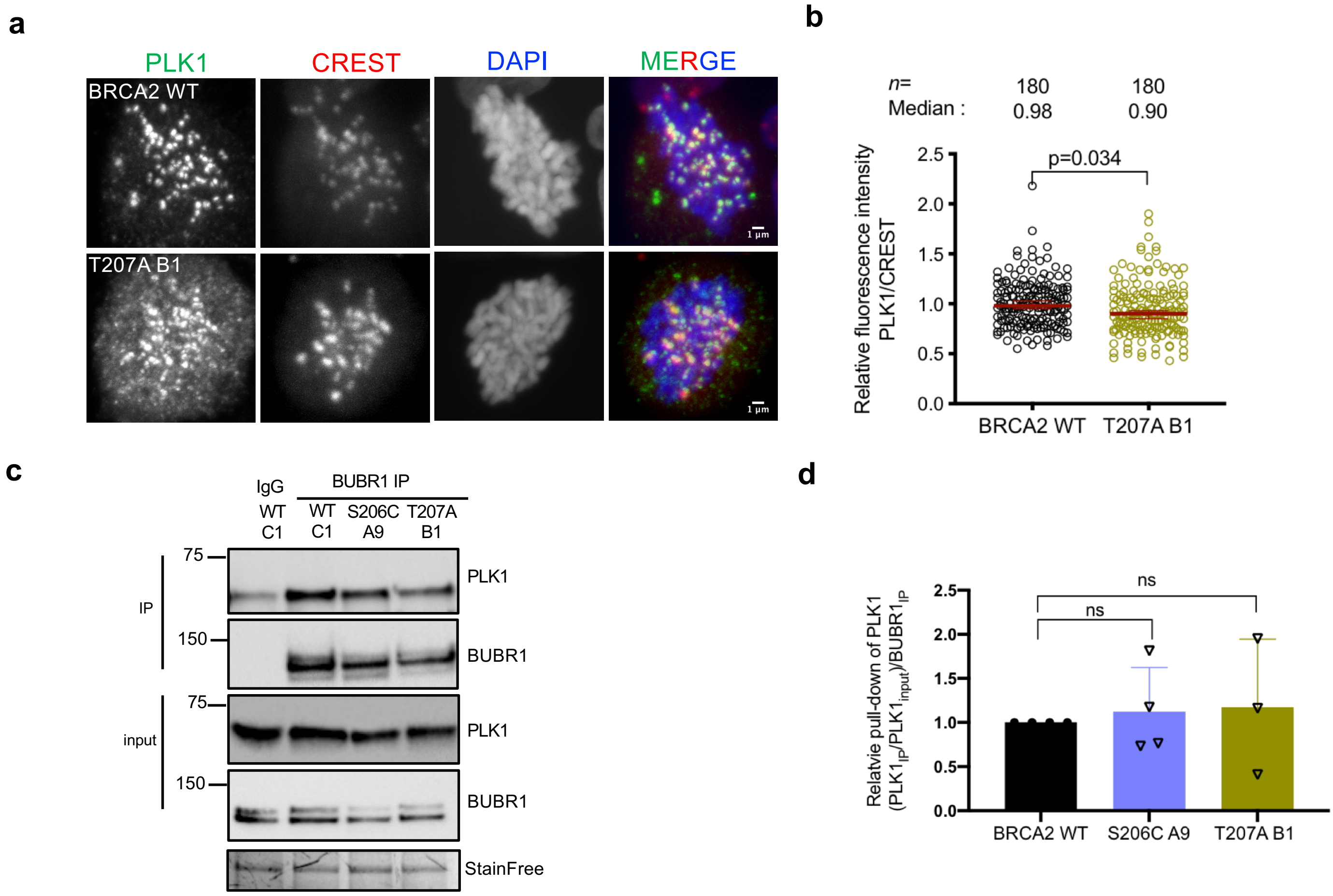

### Supplementary Fig. 7

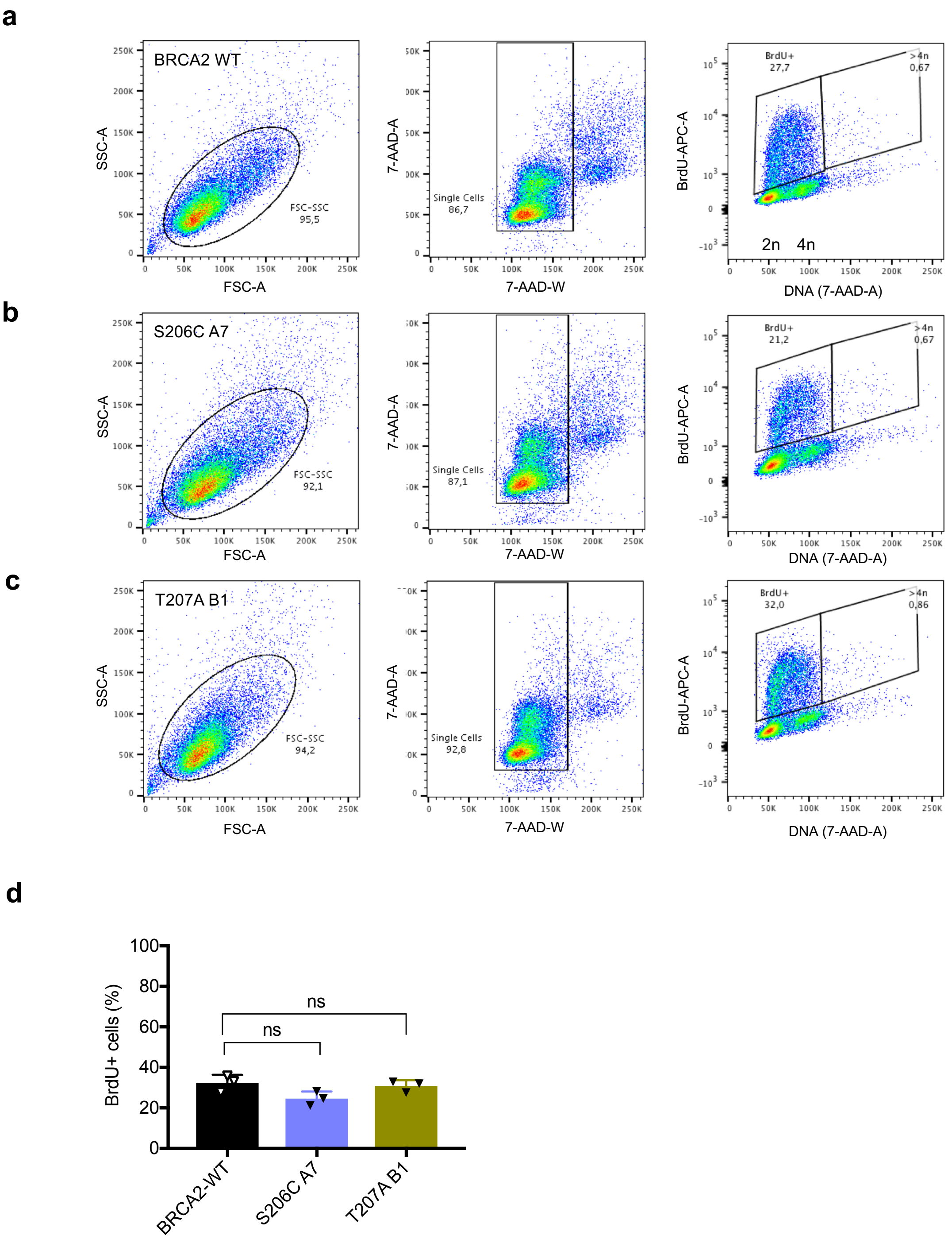

Supplementary Fig. 8

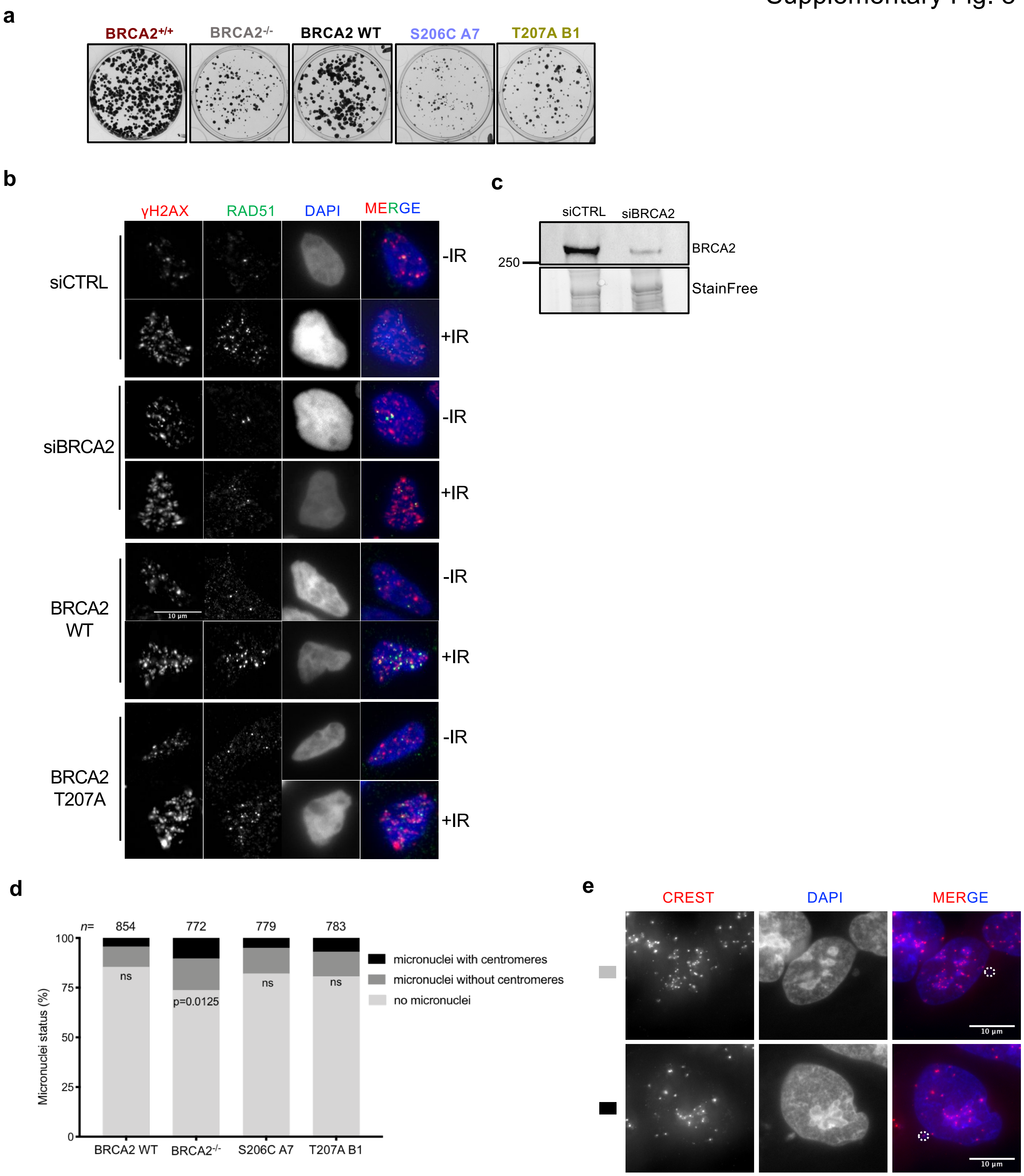

Supplementary Fig. 9

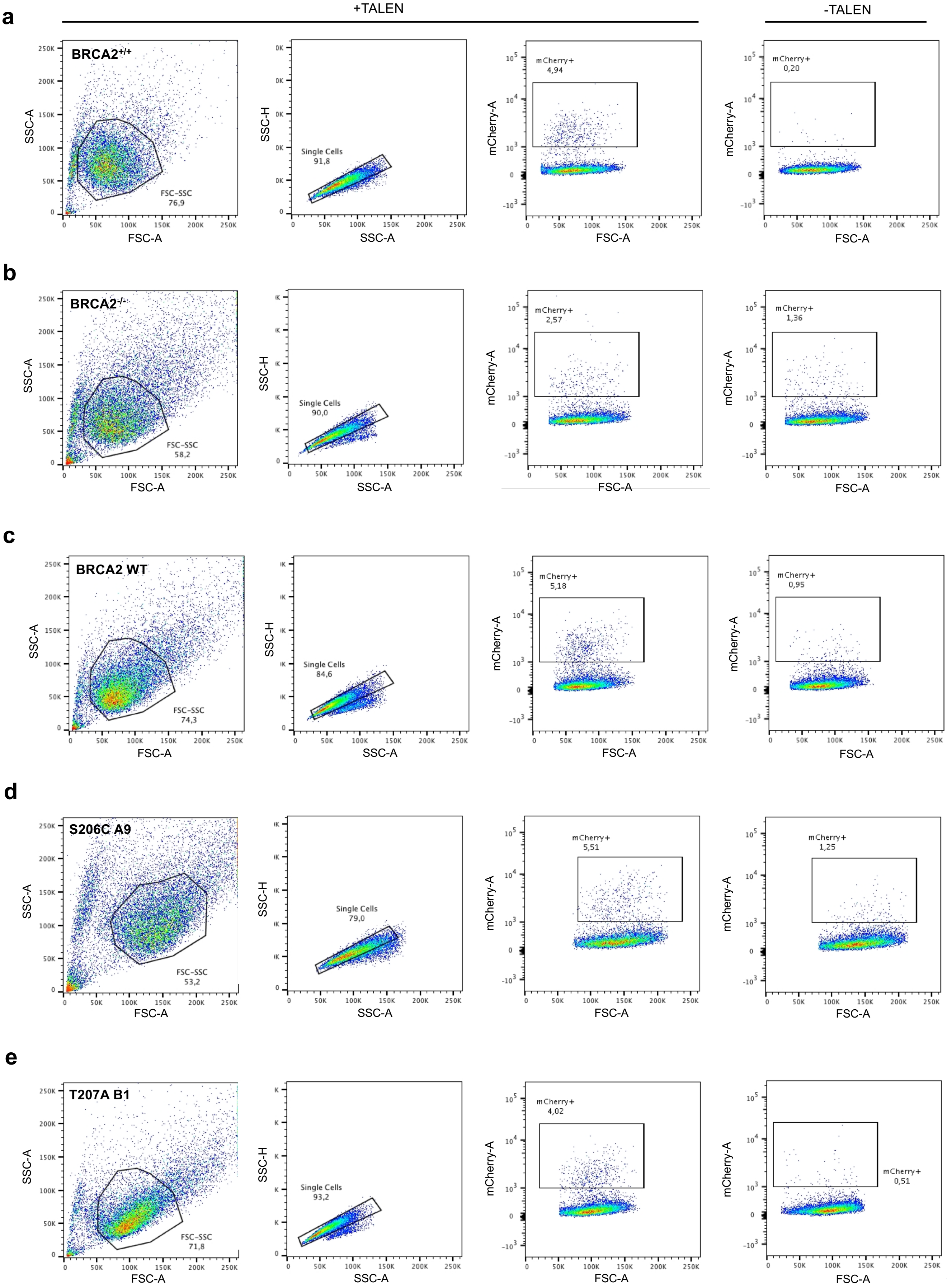

### Supplementary Figure 10

**A**

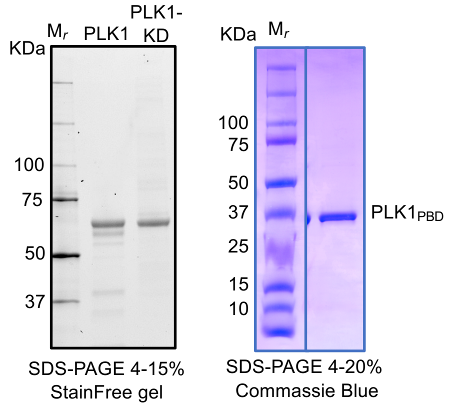


**B**

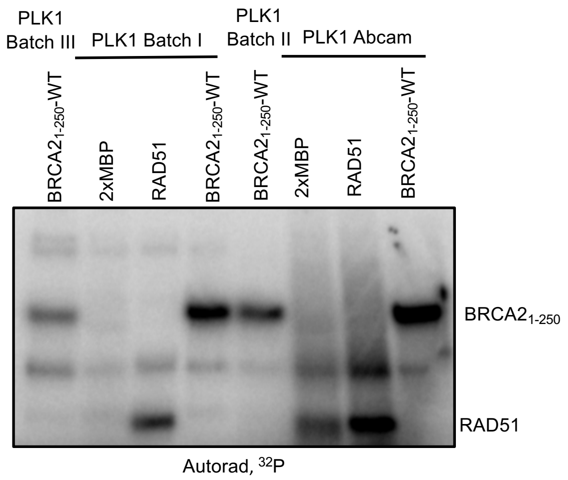
